# Modelling dementia in *Drosophila* uncovers shared and specific targets of TDP-43 proteinopathy across ALS and FTD relevant circuits

**DOI:** 10.1101/2022.11.16.516763

**Authors:** R Keating Godfrey, Reed T Bjork, Eric Alsop, Brijesh S Chauhan, Hillary C Ruvalcaba, Jerry Antone, Allison F Michael, Christi Williams, Grace Hala’ufia, Alexander D Blythe, Megan Hall, Kendall Van Keuren-Jensen, Daniela C Zarnescu

**Affiliations:** Department of Molecular and Cellular Biology, 1007 E. Lowell St, Life Sciences South, University of Arizona, Tucson AZ 85721, USA; current address: McGuire Center for Lepidoptera and Biodiversity, Florida Museum of Natural History, 3215 Hull Road, University of Florida, Gainesville, FL 32611, USA; Translational Genomics Research Institute, 445 N 5^th^ St, Phoenix, AZ 85004, USA; current address: Cellular and Molecular Physiology, 500 University Drive Crescent Building C4605, Penn State College of Medicine, Hershey PA 17033, USA

## Abstract

Amyotrophic lateral sclerosis (ALS) and fronto-temporal dementia (FTD) comprise a spectrum of neurodegenerative diseases linked to TDP-43 proteinopathy, which at the cellular level, is characterized by loss of nuclear TDP-43 and accumulation of cytoplasmic TDP-43 puncta that ultimately cause RNA processing defects including dysregulation of splicing, mRNA transport and translation. Complementing our previous models of ALS, here we report a novel model of FTD based on overexpression of TDP-43 in the *Drosophila* mushroom body (MB) circuit. This model recapitulates several aspects of FTD pathology including age-dependent neuronal loss, and nuclear depletion and cytoplasmic accumulation of TDP-43, accompanied by behavioral deficits in working memory and sleep that occur before axonal degeneration ensues. RNA immunoprecipitations identify several candidate mRNA targets of TDP-43 in MBs, some of which are unique to the MB circuit while others are shared with motor neurons. Among the latter is the glypican Dally-like-protein (Dlp), a modulator of Wg/Wnt signaling. Using genetic interactions we show that overexpression of Dlp in MBs mitigates TDP-43 dependent working memory deficits. These results highlight the utility of modelling TDP-43 proteinopathy in *Drosophila* and provide a novel platform for studying the molecular mechanisms underlying FTD, and potentially uncovering shared and circuit specific vulnerabilities in ALS/FTD.

## INTRODUCTION

Characterized by extensive overlap in cellular pathology, genetic mutations, and molecular markers, amyotrophic lateral sclerosis (ALS) and frontotemporal dementia (FTD) comprise a neurodegenerative disease spectrum affecting multiple subsets of neurons in the spinal cord, motor cortex, cingulate cortex, and frontal and temporal lobes (Arai et al., 2006; Ferrari et al., 2011; Ling et al., 2013; Neumann et al., 2006). Notably, nearly 97% of ALS cases (Mackenzie et al., 2007) and approximately 45% of FTD cases (Cairns et al., 2007) exhibit neurodegeneration with ubiquitin-positive cytoplasmic inclusions that contain transactive response (TAR) DNA binding protein 43 (TDP-43). A number of point mutations in *TARDBP*, the gene encoding TDP-43, have been identified as causative of ALS (Kabashi et al., 2008; Rutherford et al., 2008; Sreedharan et al., 2008) while others have been linked to FTD (Borroni et al., 2009; Chen et al., 2019). Although mutations in at least 20 other genes have been associated with ALS/FTD (Al-Chalabi et al., 2012), the majority of ALS/FTD cases are sporadic.

Substantiating its significance as a marker of pathology, TDP-43 has also been identified as a component of cytoplasmic aggregates in dementias other than FTD, including Alzheimer’s disease (AD) (Chang et al., 2015; Josephs et al., 2014b; Rohn, 2008), AD with Lewy body dementia (AD/LBD) (McAleese et al., 2017), Parkinsonism-dementia complex (PDC) (Hasegawa et al., 2007), and hippocampal sclerosis (Amador-Ortiz et al., 2007). Up to 57% of AD cases exhibit TDP-43 positive intracellular inclusions (Josephs et al., 2014a); notably, AD patients with TDP-43 proteinopathy show accelerated disease progression with more severe cognitive impairment (Josephs *et al*., 2014b; Meneses et al., 2021). In addition to the presence of TDP-43 proteinopathy, dementias such as FTD and AD also share TDP-43 associated cellular pathology, including dystrophic neurites (Mackenzie et al., 2011). Thus, despite differences in underlying genetic risk factors and specific neural populations affected, a large proportion of dementias exhibit overlapping TDP-43 pathology, providing a common entry point for understanding shared mechanisms of neurodegeneration (Gao et al., 2018; Jo et al., 2020).

Wild-type TDP-43 is localized primarily to the nucleus where it regulates transcription (Swain et al., 2016) and splicing (Fiesel et al., 2012; Freibaum et al., 2010; Ling et al., 2015; Polymenidou et al., 2011; Tollervey et al., 2011) necessary for nervous system development and function. TDP-43 also shuttles between the nucleus and cytoplasm and has been shown to play a role in axonal and dendritic mRNA localization (Alami et al., 2014; Chou et al., 2018; Fallini et al., 2012), stress granule dynamics (Colombrita et al., 2009; Khalfallah et al., 2018; McDonald et al., 2011) and translation (Altman et al., 2021; Coyne et al., 2017; Lehmkuhl et al., 2021). During severe or prolonged cellular stress and in disease, TDP-43 exits the nucleus and associates with ubiquitin-positive cytoplasmic inclusions (Ramaswami et al., 2013). Indeed, cellular dysfunction and subsequent degeneration associated with TDP-43 pathology have been attributed to both loss-of-function of nuclear TDP-43 (Ling et al., 2015) and cytoplasmic gain-of-function (Walker et al., 2015) whereby cytoplasmic inclusions sequester mRNAs and perturb protein homeostasis (Ramaswami et al., 2013).

Following the identification of TDP-43 as a protein of pathological interest in numerous neurodegenerative diseases, model organisms have been critical to understanding the normal biological functions of TDP-43 and how disruption of these functions leads to disease. Despite having diverged from a common ancestor with vertebrates over 600 million years ago (Erwin, 2015), the fruit fly (*Drosophila melanogaster*) genome contains a homolog of *TARDBP*, namely *TBPH*, which displays sequence similarity and functional overlap with human TDP-43 (Ayala et al., 2005; Estes et al., 2011).

In addition to extensive overlap in neurogenetic homology, circuit-level conservation in the central nervous system has also been demonstrated between invertebrates and vertebrates (Bier, 2005; Bridi et al., 2020; Strausfeld and Hirth, 2013), making invertebrate models relevant for understanding diseases of cognition. Indeed, flies display several behaviors affected in human disease (Muqit and Feany, 2002) including associative learning (Heisenberg, 2003), long-term memory (Isabel et al., 2004), sleep (Shaw et al., 2000), social interactions (Baier et al., 2002; Kamyshev et al., 2002), and addiction (Bellen, 1998) that can serve as system-level readouts of molecular- and/or cellular-level dysfunction. Models of TDP-43 proteinopathy in *Drosophila* have recapitulated the locomotor phenotypes, motor neuron dysfunction, and shortened lifespan (Estes et al., 2011; Li et al., 2010; Romano et al., 2012) observed in ALS and highlighted degeneration in central brain structures (Li et al., 2010) reminiscent of FTD. Beyond simply simulating disease states, these fly models have identified translational targets (Coyne et al., 2017; Coyne et al., 2014; Lehmkuhl et al., 2021) and genetic modifiers of TDP-43 proteinopathy (Azpurua et al., 2021; Coyne et al., 2015; Sreedharan et al., 2015; Zhan et al., 2013), and exposed systemic effects on metabolic pathways (Loganathan et al., 2022; Manzo et al., 2019).

While TDP-43 proteinopathy has been extensively studied in motor neurons and glia as a model of ALS, a robust fly model of dementia based on TDP-43 proteinopathy has not yet been established. This is in part because the mushroom bodies (MBs) of the fly brain, which were originally described as essential to odor learning (de Belle and Heisenberg, 1994) have only recently been recognized as being required for multi-modal and context-dependent learning (Liu et al., 1999; Vogt et al., 2014) including place learning (Ostrowski et al., 2015). The MBs also regulate sleep (Driscoll et al., 2021; Joiner et al., 2006), satiety (Tsao et al., 2018), social behavior (Sun et al., 2020) and gate behaviors attributed to other brain regions such as decision-making (Zhang et al., 2007) and aggression (Zwarts et al., 2015). Therefore, the MBs share functional overlap with regions of the vertebrate cortex (Wolff and Strausfeld, 2016). Existing fly models of intellectual disability (Mariano et al., 2020) and AD(Chakraborty et al., 2011) have either focused on, or identified phenotypes in the MBs of the fly brain, suggesting that this structure is appropriate for modelling diseases of cognition.

Here we describe the development of a novel fly model of FTD based on TDP-43 proteinopathy induced by overexpression of wild-type or mutant human TDP-43 in a subset of MB neurons. Our model recapitulates key cellular and behavioral characteristics of human dementias with TDP-43 pathology, including age-dependent loss of nuclear TDP-43, axonal degeneration, and working memory and sleep deficits that parallel those observed in dementia patients. Using RNA immunoprecipitations we show that in MBs, TDP-43 associates with several mRNAs, a subset of which are unique to MBs while others are shared with mRNA targets previously identified in motor neurons. Interestingly, among the latter group we identified *dlp* mRNA, encoding the heparan sulfate proteoglycan (HSPG) Dally-like protein (Dlp)/GPC6, which we previously showed that is a target of TDP-43 in fly models of ALS and altered in ALS spinal cords (Lehmkuhl et al., 2021). We report that *dlp* mRNA is enriched in TDP-43 complexes in MBs and Dlp protein is decreased within axons in an age-dependent manner, consistent with TDP-43 dependent axonal transport and/or translation deficits. Notably, Dlp overexpression in MBs rescues TDP-43 dependent deficits in working memory consistent with Dlp being a physiologically significant target of TDP-43 in the MB circuit. These findings demonstrate that TDP-43 proteinopathy in MBs causes dementia-like phenotypes that are mediated at least in part by the heparan sulfate proteoglycan (HSPG) Dally-like protein (Dlp)/GPC6, a regulator of the Wg/Wnt signaling pathway. Furthermore, we identify candidate targets of TDP-43 unique to MBs that suggest distinct neuronal vulnerabilities across the ALS – FTD spectrum.

## RESULTS

### Overexpression of TDP-43 in Kenyon cells results in age-dependent cell loss and nuclear to cytoplasmic mislocalization

The insect MBs are comprised of large, complex neurons called Kenyon cells, referred herein as MB neurons (MBNs), which form a layered network with reticular feedback (Crittenden, 1998), structural characteristics reminiscent of vertebrate cortical neurons and networks. There are three subtypes of MBNs based on developmental origin, cell morphology, and wiring. Axons of these three types form five layered lobes, the α/β, α’/β’, and γ lobes, which are further compartmentalized into columnar-like regions based on neuromodulatory inputs and excitatory or inhibitory postsynaptic partners (Aso et al., 2014). To model FTD-like TDP-43 proteinopathy we used a split-GAL4 driver line *SS01276* available from the Janelia FlyLight split-GAL4 driver collection (Aso, 2021) to express human TDP-43 (Estes et al., 2013) in a subset of MBNs that form the α/β and γ lobes of the adult *Drosophila* mushroom bodies. Mushroom body development is well-characterized in *Drosophila*. Neurons forming the γ and α’/β’ lobes are embryonic and larval in origin, respectively (Kunz et al., 2012) and persist through metamorphosis, whereas those comprising the α/β lobes are born after puparium formation (Lee et al., 1999).

We characterized expression driven by *SS01276* in the 3^rd^ instar larva (L3) and found that cytoplasmic YFP, YFP tagged TDP-43 (*TDP-43::YFP*) and membrane targeted RFP (*mCD8:: RFP*) are limited to MBNs (Figure 1-supplement 1). Signal is detected in approximately 50 larval MBNs in both TDP-43^WT^ overexpression (OE) and TDP-43^G298S^ OE brains (Figure 1A,1F *TDP-43^WT^::YFP mCD8::RFP*; Figure 1-supplement 2,*TDP-43^G298S^::YFP mCD8::RFP*). In L3, the MBs comprise approximately 60 γ lobe MBNs and 40 α’/β’ lobe MBNs (Lee et al., 1999) thus at this stage *SS01276* could possibly be driving expression in some combination of γ lobe or α’/β’ lobe MBNs. However, since adult expression of *SS01276* was determined to be limited to γ and α/β lobe MBNs (Aso, 2021), and given that both γ and α’/β’ lobe MBNs persist through metamorphosis, it is likely that the expression we observed during L3 for *SS01276* corresponds to γ lobe MBNs.

**Figure 1.**
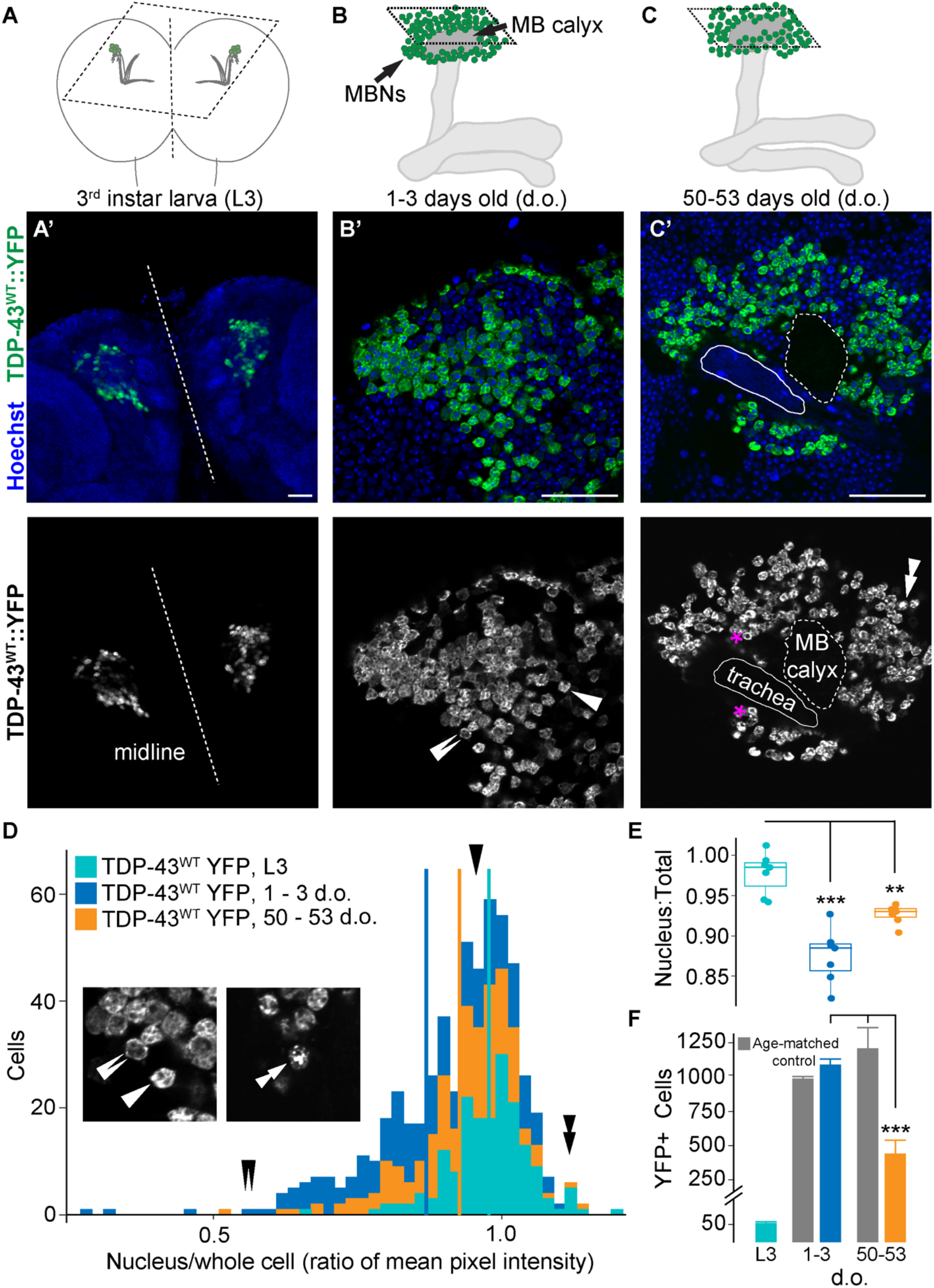
*TDP-43^WT^* overexpression in Kenyon Cells results in cytoplasmic mis-localization and age-dependent cell loss. Illustrations in (A), (B), and (C) correspond to approximate plane of imaging for Kenyon cells in (A’) larval, (B’) young adult, and (C’) old adult brain. (A’) Nuclear localization of human TDP-43^WT^ YFP in the larval MBNs. (B’) Young adult flies (1 – 3 days) contain cells with TDP-43^WT^ YFP in the cell body and cells showing nuclear loss of TDP-43^WT^ YFP (adjacent, double arrows as compared with single arrow). (C’) Old adults (50 – 55 days) have fewer cells with nuclear depletion but concurrent loss in total number of TDP-43^WT^ positive neurons (quantified in E and F). Stacked, double arrows indicate a cell that has a signal intensity ratio > 1.0 due to punctate TDP-43 signal in the nucleus. Magenta asterisks indicate cells where nuclear depletion can be detected visually. (D) Histograms showing distribution of TDP-43^WT^ YFP signal intensity ratios (cell nuclei to total cell) shift with age. Arrows correspond to measurements taken from cells shown in figure inset taken from (B’), (C’). (E) Age-specific TDP-43^WT^ YFP signal intensity ratios. (F) Decrease in TDP-43^WT^ YFP positive cells from young to old adults. *SS01276* > *YFP* used as age-matched controls. Larval cell numbers provided for reference, but not used in statistical comparisons. Data for TDP-43^G298S^ YFP presented in Figure 1-supplement 2. Scale bars = 50 μm. ** = P < 0.01; *** = P < 0.001.

Visual inspection of larval MBNs indicated that TDP-43 YFP was largely confined to the nucleus in L3 brains (Figure 1A) but became mislocalized outside the nucleus in young adults. The nucleus comprises nearly all of the cell body in MBNs, making it challenging to accurately assess cytoplasmic mislocalization or nuclear depletion. Therefore, to determine age-dependent changes in TDP-43 localization, we measured mean signal intensity of TDP-43 YFP in the nucleus by tracing the nucleus in the Hoechst channel, then measured mean signal intensity of TDP-43 YFP in the entire cell by tracing the YFP signal of the entire cell boundary (see Materials and Methods and Figure 1-supplement 3) and calculated the ratio of nucleus to total cell signal mean intensity (Figure 1D, E. Since the MBN cytoplasms are extremely thin, this ratio should be nearly 1.0 when TDP-43 is localized to both the nucleus and the cytoplasm (single arrow, Figures 1B, D, less than 1.0 when TDP-43 YFP is depleted from the nucleus and mislocalized to the cytoplasm (see double arrows, Figures 1B, D and slightly above 1.0 when TDP-43 is contained largely in the nucleus (see stacked arrows, Figures 1C, D.

Using this measure of nuclear depletion, we found a significant decrease in the mean signal intensity ratio across the population of MBNs from juvenile to young adults in the context of both TDP-43 OE variants (Figure 1E, *TDP-43^WT^::YFP mCD8::RFP* 0.978±0.005 in L3 vs. 0.869±0.009 in young adults, P < 0.0001; Figure 1-supplement 2, *TDP-43^G298S^:: YFP mCD8::RFP* 0.967±0.002 in L3 vs 0.906±0.012 in young adults, P = 0.0004; supplemental table S1E). Surprisingly, the ratio of nucleus to total cell signal mean intensity increased in old adult brains compared to young adults, however, it was still significantly lower than at the juvenile age time point (L3) in our TDP-43^WT^ OE model (Figure 1E, 0.978±0.005 in L3 vs. 0.927±0.009 in old adults, P = 0.0041; supplemental table S1E). A plausible explanation for this result is provided by our findings that there are significantly fewer TDP-43 YFP positive MBNs in old compared to younger TDP-43 OE brains or to old YFP controls (*TDP-43^WT^:: YFP mCD8::RFP,* 445±96 cells in old adults vs 1089±40 cells in young adults vs. 1206±150 cells in YFP old adults, P < 0.001; Figure 1F). Similar results were obtained for TDP^G298S^ YFP expressing brains (*TDP-43^G298S^:: YFP mCD8::RFP* 473±79 cells in old adults vs 1097±55 cells in young adults vs 1206±150 cells in YFP old adults, P < 0.001; Figure1-supplement 2; supplemental table S1F). These results are consistent with TDP-43 OE causing age-dependent cell loss and suggest that the MBNs exhibiting increased levels of cytoplasmic TDP-43 are more vulnerable and lost during aging, likely through increased cell death.

### Mushroom body axons exhibit age-dependent TDP-43 puncta accumulation and progressive degeneration

To determine the effect of TDP-43 proteinopathy on axonal integrity and overall lobe morphology, we co-expressed TDP-43 and the membrane marker mCD8 RFP. Signal intensity of mCD8 RFP showed progressive decrease in the γ lobe over time (young, 1.01±0.04, middle-aged, 0.87±0.25, and old, 0.40±0.10 fluorescence intensity ratio to age matched controls; Figure 2A, B, insets b, e, h; quantified in C; supplemental table S2A), suggesting that axonal thinning occurred in an age-dependent manner. Simultaneously, TDP-43 formed increasingly larger axonal puncta in the γ lobe with age (young, 1.00±0.04, middle-aged, 1.19±0.05, and old, 1.35±0.05 area normalized to age matched controls Figure 2B, insets c, f, i; quantified in D; supplemental table S2B). These findings are consistent with the formation of dystrophic axons, membrane fragmentation and cytoplasmic aggregation of TDP-43. A similar pattern of axonal loss and fragmentation in the γ lobe, coincident with increased TDP-43 puncta size, was observed in TDP-43^G298S^ OE flies (Figure 2-supplement 1A, B insets c, f, i, quantified in C and D; supplemental tables S2A, S2B). We could not detect statistically significant changes in mCD8 RFP signal intensity nor TDP-43 cytoplasmic puncta in the α/μ lobes of TDP-43^WT^ OE flies, however these lobes showed dramatic, age-dependent thinning not observed in controls (Figure 2B inset i). In TDP-43^G298S^ OE brains, we observed both an increase in the size of TDP-43 YFP cytoplasmic puncta within α/μ lobes between young and middle-aged adults, and a thinning of these lobes (Figure 2-supplement 1B inset i). Interestingly, these phenotypes are reminiscent of the age-dependent pathophysiology observed in human patients. Given that γ lobe neurons are born earlier than those within the α/μ lobes, the more severe phenotypes observed in this neuronal subpopulation highlight the potential contribution of aging to TDP-43 proteinopathy.

**Figure 2.**
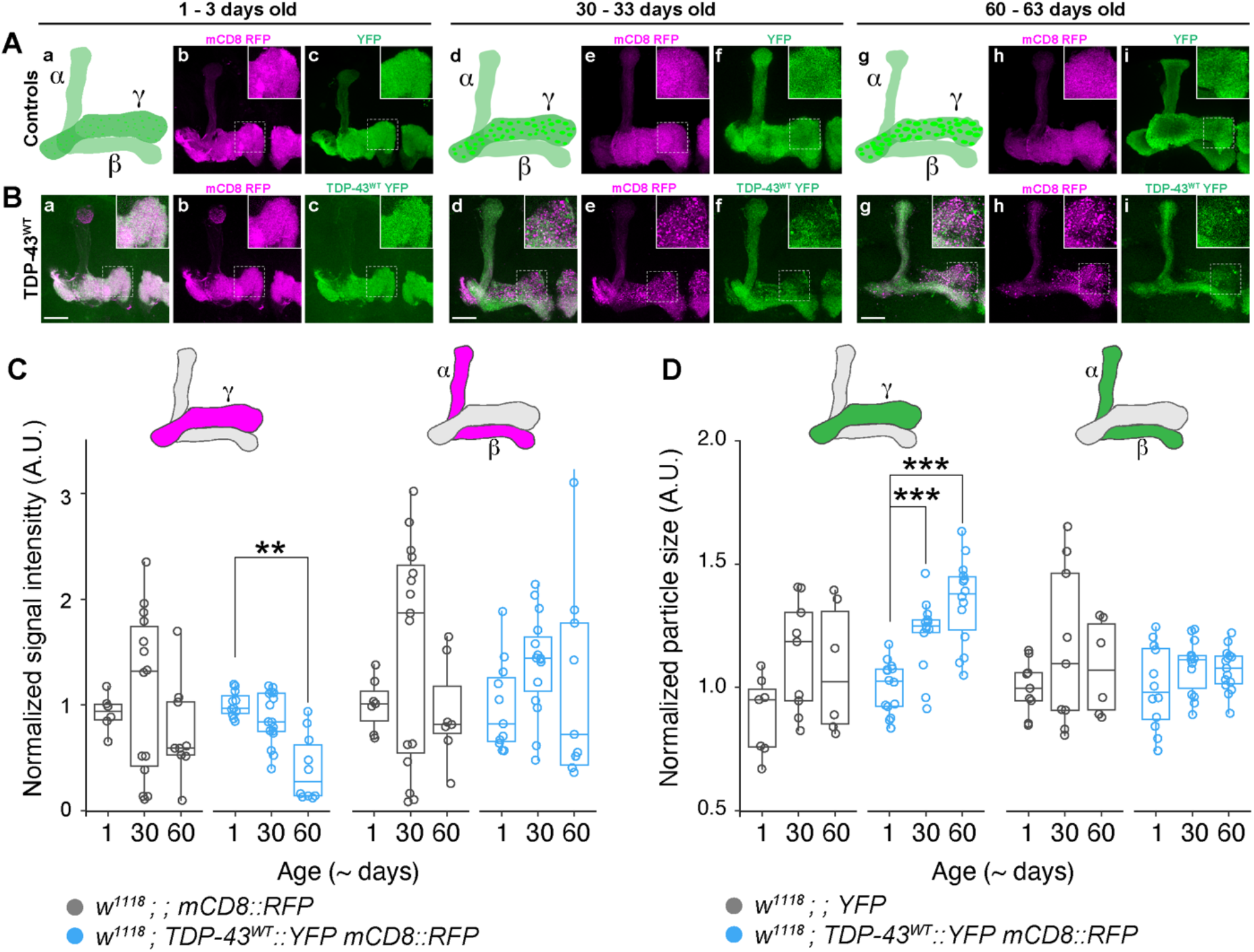
Mushroom body lobes (MBLs) show age-related, region-specific *TDP-43^WT^* accumulation and axonal fragmentation. (A) Illustrations of MBLs targeted by *TDP-43^WT^*OE and observed changes with age (a, intact in young flies; d, increasingly punctate TDP-43 YFP by middle-age; g, axonal thinning and evidence of degeneration specifically in the γ lobe). These are presented alongside morphology of young (b, c,) middle-aged (e, f) and old (h, i) control flies expressing membrane-bound RFP (mCD8 RFP) or cytoplasmic YFP. (B) Overexpression of TDP-43^WT^ YFP in MBNs results in axonal localization of TDP-43 in young adult flies (c) and dystrophic neurites in middle-aged (e) and old (h) flies. (C) Signal intensity of mCD8 RFP with age in γ (left) and α/μ (right) lobes. (D) Changes in TDP-43^WT^ YFP particle size with age in γ (left) and α/μ (right) lobes. Data for TDP-43^G298S^ presented in Figure 2-supplement 1. Scale bar = 25 μm * = P <0.05; ** = P < 0.01; *** = P < 0.001.

### TDP-43 proteinopathy causes working memory deficits

In humans, working memory is disrupted in FTD and other TDP-43 associated dementias (Poos et al., 2018; Stopford et al., 2012). To test whether flies overexpressing TDP-43 exhibit working memory deficits, we used miniature Y-mazes to measure spontaneous alternation behavior by scoring the number of three consecutive arm entries over a period of ten minutes (see Materials and Methods, Figure 3A, Lewis et al., 2017). These experiments showed that although TDP-43 OE results in increased overall movement (males: *TDP-43^WT^::YFP* in an *Oregon-R background*, 146.00±7.51 mm vs. *Oregon-R*, 106.19±8.18 mm, P = 0.018; females: *TDP-43^WT^::YFP* in an *Oregon-R background*, 110.53±7.51 mm vs. *Oregon-R*, 81.12±5.24 mm; Figure 3B; for means of individual replicates see supplemental table S3), percent alternation is significantly decreased in young adult males (*TDP-43^WT^::YFP* in an *Oregon-R background*, 0.57±0.01 vs. *Oregon-R*, 0.61±0.01, P = 0.012) and females (*TDP-43^WT^::YFP* in an *Oregon-R background*, 0.60±0.02 vs. *Oregon-R*, 0.65±0.01, P = 0.006; Figure 3C), consistent with TDP-43 proteinopathy causing working memory deficits in the MB circuit.

**Figure 3.**
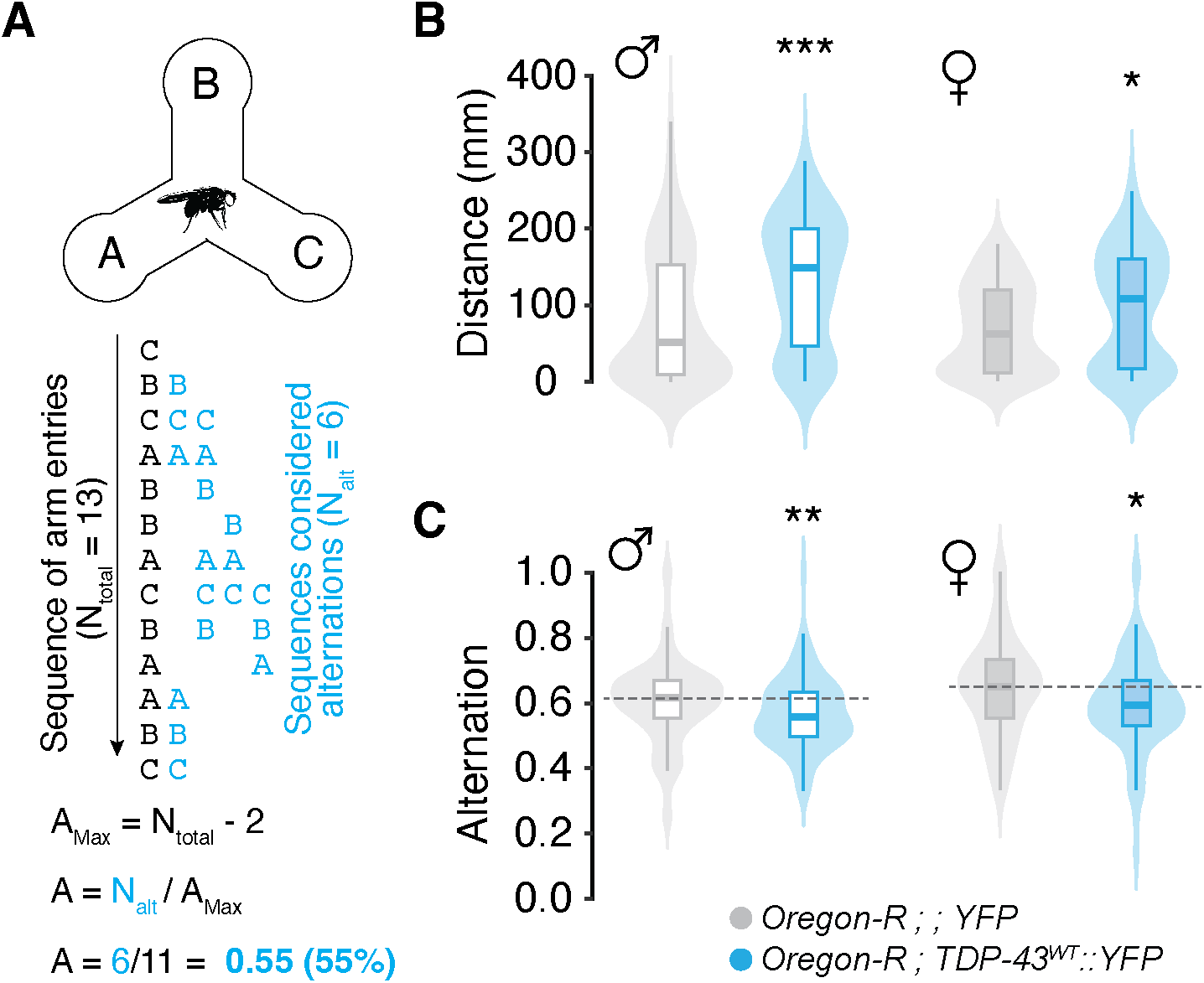
Overexpression of TDP-43 in mushroom body MBNs results in spatial working memory deficits. (A) Schematic of the spatial working memory assay showing the sequence of decisions made by a fly (black text) and those coded as alternations (blue text). Alternation scores were calculated as the number of alternations (N_alt_) divided by alternation attempts (A_max_). (B) Young (1-3 days old) flies with TDP-43^WT^ YFP OE show significant increases in movement. (C) Despite moving more, these flies show reduced alternation percent when compared with controls. Grey dotted line = control group median. A% = alternation percent. * = P <0.05; ** = P < 0.01; *** = P < 0.001.

### TDP-43 expression in the MB circuit causes sleep alterations

In humans, frontal and temporal lobe degeneration is associated with increased daytime somnolence (Sani et al., 2019), but overall sleep disturbances are highly variable in FTD and may be detectable in earlier stages than in AD (Bonakis et al., 2014). To determine whether TDP-43 proteinopathy in the MB circuit alters sleep, we used *Drosophila* Activity Monitors (DAMs, see Materials and Methods) to measure total daytime and nighttime sleep. Our experiments show that on average, both male and female TDP-43 OE flies appear to be lethargic; they spent a greater proportion of their time sleeping during both the day and night compared to controls (approximately 11% more for TDP-43^WT^ OE and approximately 8% more for TDP-43^G298S^ OE). However, there are subtle differences in how TDP-43 OE affects sleep patterns based on age and sex. While males consistently sleep more during the day, this effect is weaker in females flies where we found trends towards increased daytime sleep at some but not all age time points in each genotype (Figure 4A; supplemental table S4A). At night, males and females also exhibited a general tendency towards increased sleep with the exception of young TDP-43 OE adult males which, when compared to *w^1118^* controls, showed signs of sleep fragmentation as evidenced by more frequent, shorter sleep bouts (Figures 4B, C for TDP-43^WT^ OE and Figure 4-supplement 1B, C for TDP-43^G298S^ OE; supplemental tables S4C sleep bout length and S4C sleep bout number).

**Figure 4.**
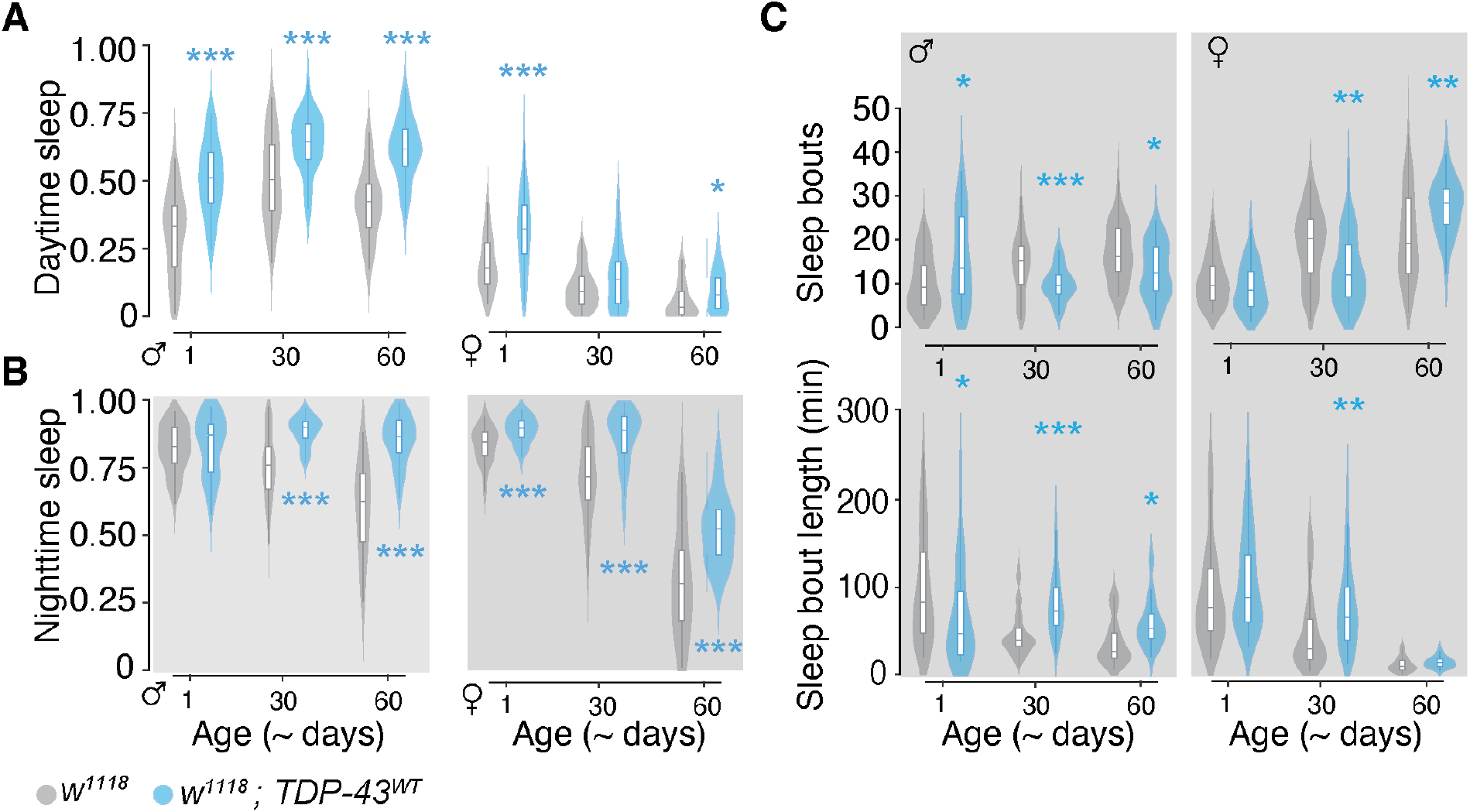
TDP-43^WT^ overexpression in mushroom body MBNs reduces arousal, increasing day and night sleep. (A) Proportion of time flies spent sleeping during the day and (B) at night. (C) Sleep fragmentation assessed by number of sleep bouts (top panel) and mean bout length (bottom panel) during the night. Male data on the left and female data on the right in each panel. Data for TDP-43^G298S^ presented in Figure 4-supplement 1. * = P <0.05; ** = P < 0.01; *** = P < 0.001

Interestingly, both humans and flies show decreased nighttime sleep with age (Li et al., 2018); in line with this, *w^1118^* flies of both sexes showed age-related declines in nighttime sleep, whereas TDP-43 OE males showed no such declines (all pairwise comparisons P > 0.05) and female declines appear less steep (TDP-43^WT^ OE, Figure 4B; TDP-43^G298S^ OE, Figure 4-supplement 1; supplemental tables S4A), consistent with our findings that TDP-43 proteinopathy causes lethargy. By middle age, TDP-43^WT^ OE males showed a pattern of longer, less frequent nighttime sleep bouts as compared with *w^1118^* (Figure 4C). Within each genotype, female w^1118^ and TDP-43^WT^ OE flies exhibited age-related sleep fragmentation as evidenced by a greater number of shorter bouts at 60 - 63 days compared with 1 – 3 days. TDP-43 OE flies still spent a greater proportion of their time sleeping than controls (Figure 4B; supplemental table S4A), an effect driven by longer sleep bouts at this aged time point (Figure 4C; supplemental table S4C). Taken together, these results indicate that TDP-43 OE in the MB circuit causes sleep deficits resembling those reported in FTD patients, including increased sleep during the day and sleep fragmentation at night, albeit the latter was detected in young males only. Notably, the sleep fragmentation phenotype occurs concomitant with the nuclear depletion of TDP-43 and before axonal degeneration suggesting that this may be a direct consequence of TDP-43 mislocalization while lethargy is likely an indirect consequence of large-scale neuronal degeneration.

### TDP-43 overexpression in Kenyon cells is sufficient to reduce lifespan in Drosophila

Life expectancy for FTD patients after diagnosis is approximately 8 – 12 years and, while patients do not often die from the disease itself, there is a measurable, if subtle, effect on life expectancy (Kansal et al., 2016; Loi et al., 2021). We therefore tested whether TDP-43 proteinopathy affects lifespan in our fly model and found that overexpression of TDP-43 in MBNs significantly reduces median survival time (TDP-43^WT^ OE, 84 days, 95% CI [84,84]; TDP-43^G298S^ OE, 89 days, 95% CI [88,91]) when compared with *YFP* controls (94 days, 95% CI [90, 95]; P < 0.0001 and P = 0.028, respectively; Figure 5 and Figure 5-supplement 1). These results indicate that overexpression of TDP-43 in a subset of MBNs in flies is sufficient to produce a nearly 10% reduction in median lifespan.

**Figure 5.**
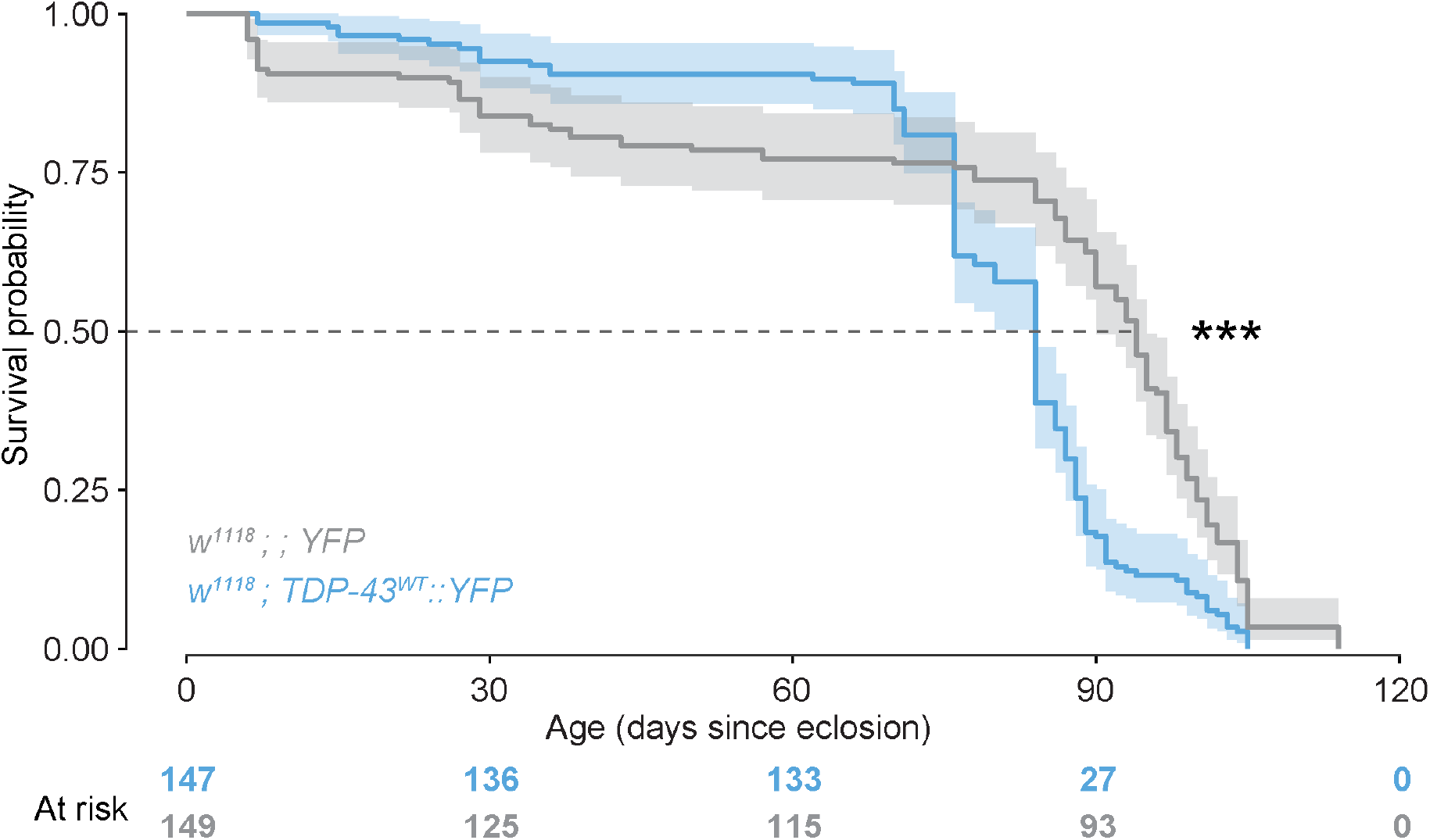
TDP-43 overexpression in MBs is sufficient to reduce lifespan. Data from male and female flies pooled for analysis. Number of flies N=149 for YFP controls and 147 for TDP-43^WT^ YFP expressing flies). Data for TDP-43^G298S^ presented in Figure 5-supplement 1. *** P < 0.001

### mRNAs enriched with TDP-43 in mushroom body neurons show partial overlap with those previously identified in motor neuron and include components of the Wnt/Wg signaling pathway

To understand how TDP-43 overexpression influences the MBN proteome, we immunoprecipitated YFP-tagged TDP-43^WT^ or TDP-43^G298S^ from *Drosophila* heads overexpressing TDP-43 YFP in MBNs using the *SS01276* Split-GAL4 driver. Next, we used DESeq2 (Love et al., 2014) to assess differential expression of mRNAs enriched in TDP-43 complexes compared to the whole brain transcriptome. We chose to identify TDP-43-enriched candidate mRNA targets using young adult flies (1 – 3 days post-eclosion), before age-dependent axonal neurodegeneration is detected in the MB lobes, to avoid detection of gene expression alterations that could be an indirect consequence of neurodegeneration. Of the 13,302 transcripts detected in the fly brain, we identified 1055 and 1393 candidate mRNA targets based on their enrichment in TDP-43^WT^ YFP and TDP-43^G298S^ YFP complexes, respectively (Log2FC > 1 compared to the brain transcriptome, P_adj_ < 0.05) (Figure 6A, Figure 6-supplement 1A). A total of 876 candidate mRNAs were shared by both variants (83% of targets in TDP-43^WT^ OE and 63% of targets in TDP-43^G298S^ OE), for a total of 1572 mRNAs enriched across both models (with 179 unique to TDP-43^WT^ OE and 517 to TDP-43^G298S^ OE, respectively). Notably, differential expression analyses also recovered previously identified TDP-43 target mRNAs including *futsch* (TDP-43^WT^ OE Log2FC = 1.81, P_adj_ = 2.66E-07; TDP-43^G298S^ OE Log2FC = 2.14, P_adj_ = 9.57E-10) (Coyne et al., 2014) and *dscam2* (TDP-43^WT^ OE Log2FC = 1.29, P_adj_ = 6.01E-04; TDP-43^G298S^ OE Log2FC = 1.81, P_adj_ = 2.42E-3) (Xiao et al., 2011).

**Figure 6.**
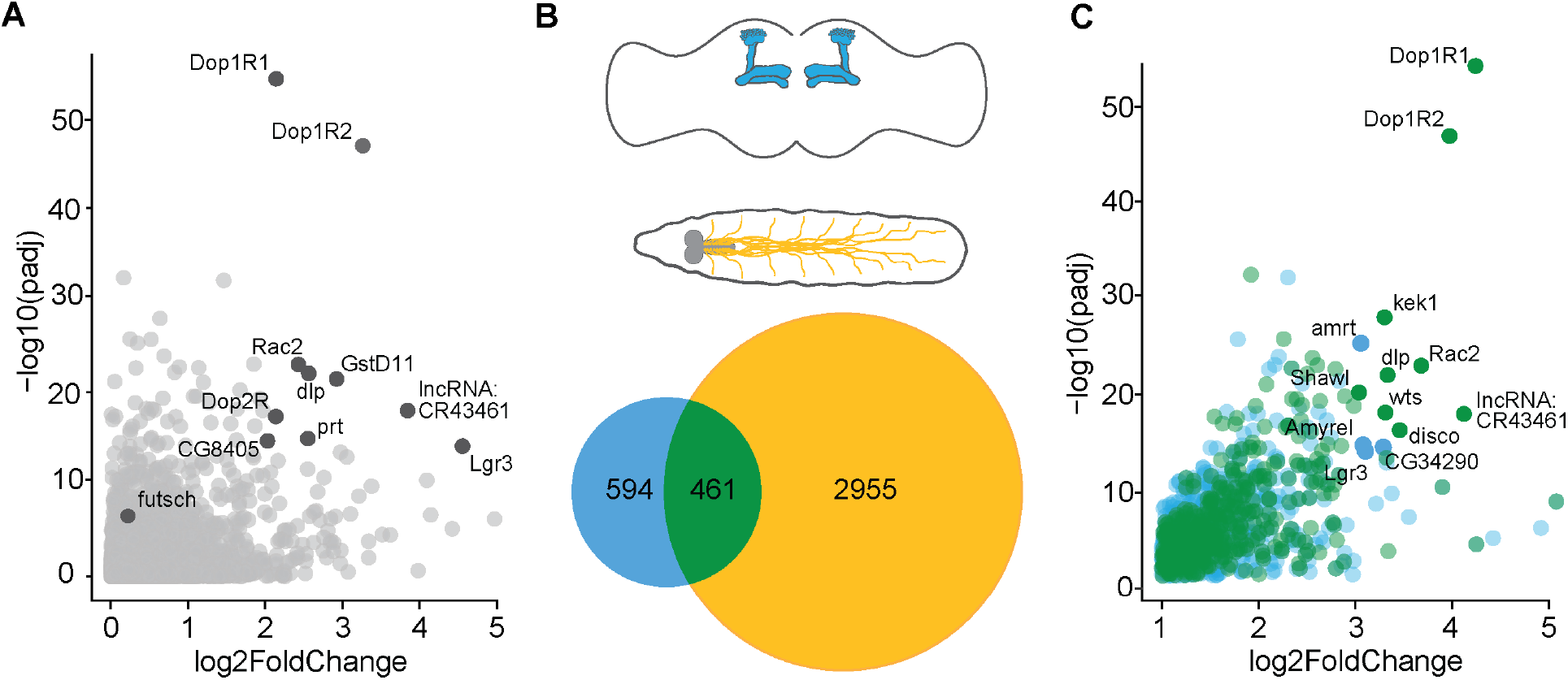
mRNAs enriched with TDP-43 in *Drosophila* models of proteinopathy in FTD and ALS. (A) Volcano plot displaying mRNAs enriched with TDP-43. Y-axis depicts Log2 Fold Change after subtraction of YFP control values (see Materials and Methods). A small number of targets showing greatest enrichment with high confidence are highlighted with blue circles (log2 Fold Change > 2 and P < 1 x 10^-14^). (B) Schematic showing subsets of neurons in fly models of FTD (MBs, above) and ALS (MNs, below), and Venn diagram depicting enriched mRNAs unique to MB and MN circuits, and overlapping between models. (C) Volcano plot displaying mRNAs enriched in TDP-43 complexes that are MB-specific (blue) or shared between MB and MN models (green); saturated green circles indicate shared targets that show log2 Fold Change > 3 and P < 1 x 10^-14^; blue circles indicate MB targets with log2 Fold Change > 2 and P < 1 x 10^-14^. Data for TDP-43^G298S^ presented in Figure 6-supplement 1.

While TDP-43 has been found to regulate mRNA processing in a cell type specific manner, there is also overlap among TDP-43 splicing targets in neuronal and muscle cell lines (Susnjar et al., 2022). Consistent with these findings, we found that nearly 44% of TDP-43^WT^ OE and 42% of TDP-43^G298S^ OE highly associated mRNAs in MBNs overlapped with those we identified previously in motor neurons (MNs) when modelling ALS by overexpressing TDP-43 with the D42 GAL4 driver (Figure 6B, Figure 6-figure supplement 1C).

Functional annotation of mRNA candidate targets in the MBs identified neuron-specific as well as neuromuscular junction molecular pathways in both the TDP-43^WT^ OE and TDP-43^G298S^ OE models (Figure 6-supplement 2A). Components of the Wingless (Wg/Wnt) signaling pathway were overrepresented in the TDP-43 proteinopathy model of FTD (Figure 6-supplement 2B). Highly enriched members of these pathways include *frizzled* (*fz*), a Wg/Wnt receptor, *dally-like protein* (*dlp*), a heparan sulfate proteoglycan (HSPG) that interacts with multiple Wnt ligands (Waghmare et al., 2020), and the Wg/Wnt receptor *frizzled 2* (*fz2*) (Yan et al., 2009). Interestingly, *dlp* mRNA was also found to be enriched with TDP-43 in fly motor neurons, where TDP-43 proteinopathy causes depletion of Dlp from the neuromuscular junction and simultaneous accumulation of Dlp puncta in cell bodies (Lehmkuhl et al., 2021). To validate that *dlp* mRNA is a target of TDP-43 in MBNs as predicted by RNA seq, we used RNA IPs followed by RT-qPCR and confirmed that *dlp* mRNA is enriched in immunoprecipitated TDP-43^WT^ and TDP-43^G298S^ complexes compared to input (Figure 6-supplement 1).

Among the mRNAs enriched with TDP-43 in fly MBNs we identified several orthologs of human genes exhibiting altered expression in the frontal cortices of FTD patients including the Ras family member RHOQ (*Rac2*, *TDP-43^WT^*Log2FC = 3.68, P_adj_ = 9.66E-24), the glypican GPC4 (*dlp*, *TDP-43^WT^* Log2FC =3.33, P_adj_ = 9.14E-23), heat shock 70kDa protein 2, HSPA2 (*Hsp70Bb*, *TDP-43^WT^*Log2FC = 2.77, P_adj_ = 1.59E-7), and the frizzled family receptor protein FZD7 (*fz*, *TDP-43^WT^* Log2FC = 2.18. P_adj_ = 4.25E-12) (Santiago et al., 2020).

In addition to TDP-43 candidate targets shared in MB and MN circuits, we also assessed the pool of mRNAs significantly enriched with TDP-43 in MBNs only, higlighting circuit specific aspects of TDP-43 proteinopathy. For TDP-43^WT^ OE we found 461 enriched mRNAs that are shared in both MB and MN, and 594 mRNAs uniquely enriched in MBs (Figure 6). DAVID analyses of the unique candidate targets for TDP-43^WT^ OE in MBs uncovered several kegg pathways altered in the context of TDP-43 proteinopathy including “Basal transcription factors”, “RNA polymerase”, “Ribosome biogenesis” and “Spliceosome” (supplemental table S6A). For TDP-43^G298S^ we found 592 enriched mRNAs that are shared in both MB and MN, and 802 mRNAs uniquely enriched in MBs (Figure 6-supplement 1). DAVID analyses of TDP-43^G298S^ specific targets in the MB circuit revealed several significantly represented pathways including “mTOR signaling pathway”, “Other types of O-glycan biosynthesis”, “TGF-beta signaling pathway”, “Nicotinate and nicotinamide signaling pathway”, “Autophagy”, “Nucleotide metabolism” and “Spliceosome” (supplemental table S6B).

Taken together, these results show that while some mRNAs enriched with TDP-43 in the MB circuit are specific to MBNs, others are shared with MNs and highlight the Wg/Wnt signaling pathway as a functional target of TDP-43 proteinopathy across the ALS/FTD spectrum.

*TDP-43 proteinopathy causes severe age-related loss of Dally-like protein in MBNs* Given *dlp* mRNA’s enrichment with TDP-43, we hypothesized that the expression of Dlp protein, a Wg/Wnt regulator, may be affected in MBNs, as we previously found in motor neurons (Lehmkuhl et al., 2021). While Dlp has been shown to be highly expressed in developing MBN axons, where it is thought to be involved in axon guidance (Rawson et al., 2005), its expression and function in adult MBNs has not been characterized. We found that, despite *dlp* mRNA being enriched in TDP-43 complexes in young adult brains, the levels and distribution of Dlp protein in TDP-43 OE flies are comparable with controls at that age (Figures 7A, B). Interestingly, Dlp levels decline with age in both controls and TDP-43 OE flies, however while Dlp is still visible in control fly brains at 50 days, Dlp is nearly undetectable in the brains of TDP-43 OE flies (Figures 7A, B, P < 0.05). A similar age-dependent reduction in Dlp expression was also detected in the context of TDP-43^G298S^ OE in MBNs (Figure 7-supplement 1). These findings are consistent with *dlp* mRNA being a target of TDP-43 in MBNs and suggest mRNA localization and/or translation as possible mechanisms for the observed reduction in Dlp protein expression.

**Figure 7.**
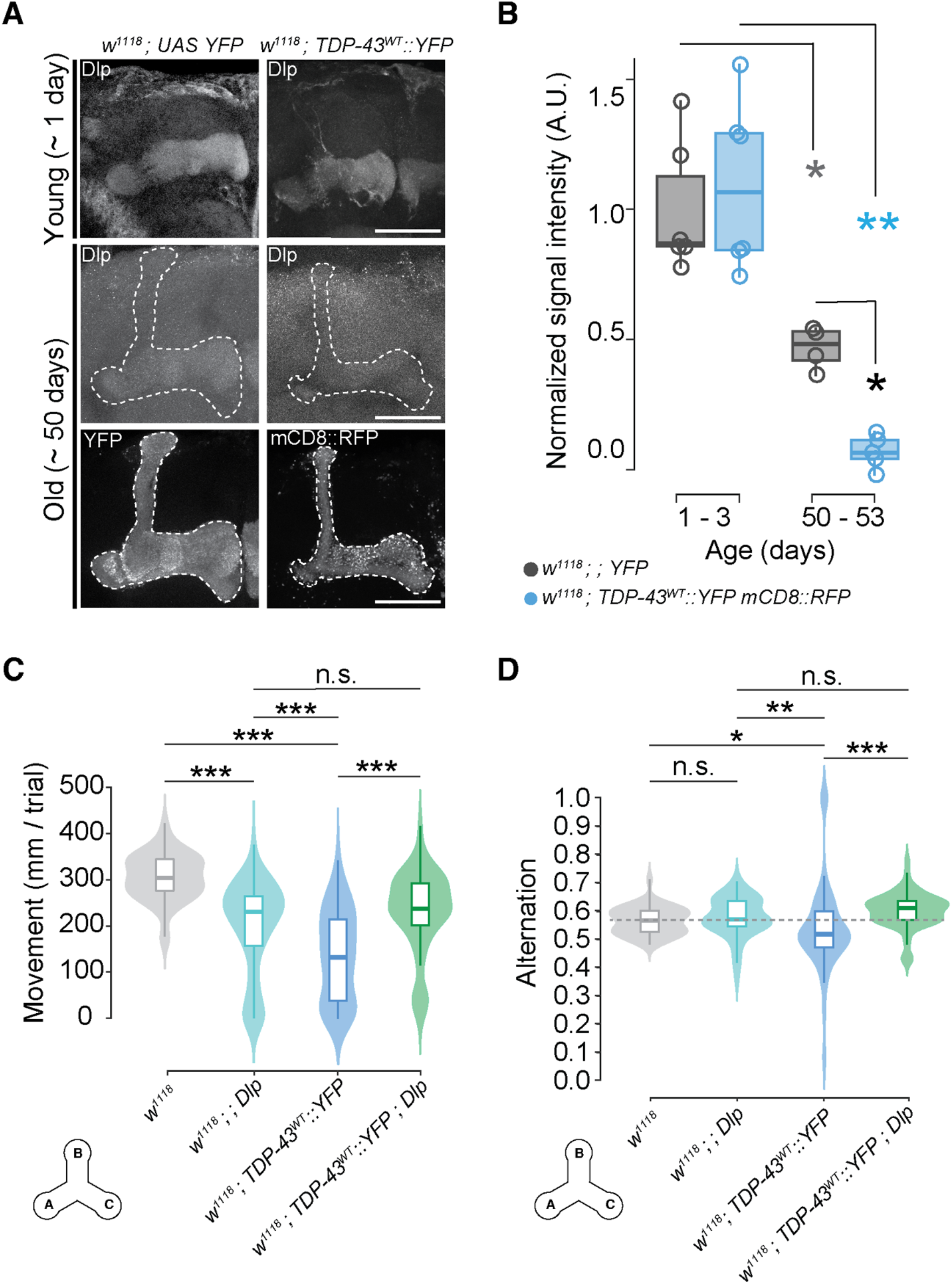
Dally-like protein is a target of TDP-43^WT^ in MBNs where it mediates TDP-43 dependent working memory deficits. (A) Dlp antibody labelling in mushroom bodies shows age and TDP-43 dependent reduction. Genotypes (*w^1118^; YFP* and *w^1118^; TDP-43^WT^::YFP mCD8::RFP*), stainings (Dlp, YFP, RFP) and ages (1 and ∼ 50 days old), as indicated. (B) Quantification of age and TDP-43 dependent changes in Dlp signal intensity within MBLs. (C) Mean distance moved and (D) alternation per trial in males overexpressing Dlp (*w^1118^ ; ; dlp OE),* TDP-43^WT^ (*w^1118^ ; TDP-43^WT^::YFP)* or both *(w^1118^ ; TDP-43^WT^::YFP; dlp OE*) compared to *w^1118^* genetic background controls. Grey dotted line represents control group median. Scale bar = 50 μm. Data for TDP-43^G298S^ presented in Figure 7-supplement 1. * = P < 0.05, ** = P < 0.01

### Dlp overexpression rescues TDP-43 dependent working memory deficits

We next evaluated the physiological significance of *dlp* mRNA as a target of TDP-43 in MBs. To address this, we tested whether overexpression of *dlp* mRNA (dlp OE) in the context of TDP-43 proteinopathy could mitigate TDP-43 dependent working memory deficits. Using the Y-maze working memory assay we found that the percent alternation deficits displayed by TDP-43^WT^ OE flies could indeed be mitigated by dlp OE in MBNs (Figure 7D). To account for altered movement caused by TDP-43 OE (*TDP-43^WT^::YFP*, 147.52 ± 13.52 mm, *w^1118^*, 229.89 ± 7.43 mm, P < 0.0001, see Figure 7C), we also compared distance-scaled alternations across genotypes and found that TDP-43 dependent alternation deficits were still mitigated by dlp OE *(TDP-43^WT^::YFP T*, 0.0221 ± 0.0028 alternation/mm, *TDP-43^WT^::YFP dlp OE*, 0.0286 ± 0.0036 alternation/mm, P < 0.0001) while no significant difference was detected between co-overexpression of TDP-43 and Dlp, and dlp OE alone (*TDP-43^WT^::YFP dlp OE*, 0.0286 ± 0.0036 alternation/mm, *dlp OE*, 0.0252 ± 0.0032 alternation/mm, P > 0.05, not shown). We note that although total movement in the Y-maze assay is influenced by the genetic background (*w^1118^* versus *OR-R*, compare Figures 3B and 7C), the significant reduction in spontaneous alternation is consistent in TDP-43 OE flies in both cases (Figures 3C and 7D). These experiments indicate that dlp OE in the MB circuit mitigates TDP-43^WT^ OE dependent working memory deficits as evidenced by the rescue of alternation behavior in the Y-maze assay. Notably, this rescue effect is not due to reduced TDP-43 expression or to diluted GAL4 activity by the presence of an additional UAS promoter, as evidenced by similar TDP-43 levels with or without dlp OE (Figures 7-supplement 2, supplement 3). Surprisingly, OE of TDP-43^G298S^ in MBNs did not cause a significant spontaneous alternation phenotype (see raw data in Supplemental Table 7). Taken together, these findings support the notion that *dlp* mRNA is a functional target of TDP-43 *in vivo* and mediates, at least in part, TDP-43 dependent, FTD relevant phenotypes in *Drosophila*.

## DISCUSSION

Neurodegenerative diseases have historically been classified by shared symptomology and post-mortem pathology. More recently, the integration of molecular markers and genetic testing has uncovered evidence that cell-type susceptibility and dysregulation of specific molecular pathways can interact to produce similar clinical presentations across different disease etiologies (Pennington et al., 2011). These findings highlight convergent pathomechanisms during the degeneration process (Gao et al., 2018; Ling et al., 2013) and suggest inroads for better understanding of these complex diseases.

A common pathological feature of several neurodegenerative diseases including ALS, FTD and other dementias is the nuclear depletion of the RNA binding protein TDP-43 accompanied by the accumulation of cytoplasmic, ubiquitinated foci (Mackenzie et al., 2010) (Amador-Ortiz et al., 2007; Chang et al., 2015; Josephs et al., 2014b; McAleese et al., 2017; Rohn, 2008). Since these dementias are often confirmed post-mortem by the presence of TDP-43 pathology, animal models play an important role in elucidating the molecular mechanisms underlying TDP-43 proteinopathy and its role in neurodegeneration. Among these, fly models have proven their utility in this research landscape in part because the availability of powerful molecular and genetic tools allowing multiple hypotheses to be pursued in parallel on a large scale. Indeed, fly models of TDP-43 proteinopathy based on loss of function or overexpression in the entire nervous system (Feiguin et al., 2009; Lu et al., 2009; Romano et al., 2012), or in subsets of neurons including associative regions of the central brain (Li et al., 2010), motoneurons (Estes et al., 2011) or the retina (Elden et al., 2010) have revealed a broad repertoire of altered pathways that have subsequently been confirmed in patient tissues (Coyne et al., 2017; Coyne et al., 2014; Lehmkuhl et al., 2021).

It has been previously shown that overexpression of TDP-43 in all MBNs using the driver line OK107 GAL4 (RRID:BDSC_854) caused age-dependent cell loss and axonal degeneration in MBs, however degeneration severity varied widely across nearly isogenic individuals, challenging its utility as a disease model (Li et al., 2010). This variability could be in part attributed to driver line “leakiness” as OK107 GAL4 drives expression beyond MBs, in optic lobe neuropils (Aso et al., 2009; Morante and Desplan, 2008) and neurosecretory cells (Enell et al., 2010; Hampel et al., 2011).To develop a robust and MB specific model of TDP-43 driven dementia in *Drosophila melanogaster,* we leveraged a split-GAL4 driver line with limited expression to a subset of Kenyon cells (γ, αβ MBNs). Notably, this model exhibits nuclear depletion and cytoplasmic accumulation of TDP-43, accompanied by age-dependent axonal degeneration and cell loss, all of which recapitulate key aspects of disease pathology. Importantly, the onset of FTD relevant behavioral symptoms is detectable prior to widespread degeneration, as observed in human disease. Additionally, this model identifies both novel, MB specific, and motor neuron-shared mRNA candidate targets that have previously been associated with TDP-43 pathology. Of the latter, here we chose to focus on the glypican Dlp, a Wg/Wnt signaling regulator and show that Dlp is a functional target of TDP-43 in the MB circuit that mediates, in part, TDP-43 dependent working memory deficits in the fly model.

Neurodegeneration in ALS/FTD is thought to be driven by both loss of nuclear function, as some nuclei become devoid of TDP-43, and cytoplasmic gain of toxic function, as evidenced by TDP-43 accumulation in cytoplasmic puncta. While thus far invertebrate models of TDP-43 proteinopathy have not faithfully recapitulated TDP-43 mislocalization (e.g., Ash et al., 2010; Estes et al., 2013), here we found that TDP-43 overexpression in MBNs more closely resembles disease pathology. Indeed, TDP-43 displays normal localization in the juvenile stage but becomes depleted from the nucleus and mislocalized to the cytoplasm in young adults.

Although changes in TDP-43 localization could reflect developmental regulation (Park et al., 2020), our finding that the number of cells with increased cytoplasmic TDP-43 is reduced from young to old age concomitant with an overall reduction in MB neuron numbers, suggests that cytoplasmic TDP-43 is toxic. Axonal degeneration is evident only in older adult flies and differentially affects MBNs that form the α/β and γ lobes, with the latter showing the greatest degeneration severity. This is particularly interesting because MBNs that form the γ lobe are embryonic in origin and therefore older than the pupal-born MBNs that form the α/β lobes (Ito et al., 1997; Kunz et al., 2012; Yu and Lee, 2007), highlighting the aging component of neurodegeneration. It is also possible that the differences in the onset and effects of TDP-43 proteinopathy we observed in γ lobe MBNs may be due to their remodelling during metamorphosis (Kunz et al., 2012), which may increase their vulnerability to TDP-43. Taken together, these findings parallel differential susceptibility observed in human cortical neurons (Nana et al., 2019; Seeley, 2008). Given the availability of split-GAL4 driver lines for subsets of MBNs (Aso et al., 2014) future studies could focus on better understanding what drives this susceptibility by modelling TDP-43 proteinopathy in various neuronal subpopulations within the MB circuit. It will be interesting to combine these powerful genetic tools with single cell RNA seq efforts in patients and flies in order to pinpoint specific cell sub-types that may be more vulnerable.

Frontotemporal lobar degeneration with TDP-43 proteinopathy (FTLD-TDP) is pathologically classified based on the cortical layers affected, the number of dystrophic neurites, and the extent of neuronal cytoplasmic inclusions (Mackenzie et al., 2011). This pathology is also commonly observed in AD/LBD that exhibit TDP-43 proteinopathy, although different brain regions are selectively affected (Josephs et al., 2014a; Josephs et al., 2014b). Interestingly, we find that neuronal cytoplasmic inclusions are absent in juveniles, appear first in young adult brains and increase in size during aging. This neuropathology precedes observations of dystrophic neurites, which are first visible around middle age, increase in severity during aging, and ultimately degenerate. This age-dependent pathology observed in the fly brain parallels that observed in human disease and provides a platform for studying the progression of neurodegeneration in the genetically tractable *Drosophila* model.

In humans, behavioral symptoms of FTD precede widespread degeneration by years suggesting that behavioral deficits arise prior to neuronal pathology, in the absence of complete loss of nuclear TDP-43 function or cytoplasmic accumulation (Perry et al., 2017). Therefore, we chose to focus primarily on working memory and sleep deficits using young adult flies, in which a subset of MBNs show nuclear depletion accompanied by widespread TDP-43 mislocalization to the axonal cytoplasm while MBN axons still appear largely intact. At this young age, we found both working memory and sleep fragmentation phenotypes, albeit the latter were only significant in males. We also assessed these behaviors in aged flies, however the distinct age-dependent deficits exhibited by the controls themselves confounded the detection of TDP-43 specific phenotypes.

Behavioral variant frontotemporal dementia (bvFTD) is diagnosed based on often subtle behavioral symptoms including a lack of empathy, increased apathy and compulsive behaviors, and deficits in executive function (Rascovsky et al., 2011). FTD was classically distinguished from AD using a perceived lack of memory deficits, however recent patient studies and meta-analyses indicate that memory deficits are common in bvFTD and at times indistinguishable between AD and bvFTD (Graham et al., 2005; Pennington et al., 2011), challenging the validity of the exclusion of episodic memory deficits in the clinical diagnoses of bvFTD (Hornberger and Piguet, 2012). Although sleep symptoms are notoriously variable and understudied across different FTD diagnoses, the most common sleep disturbances observed in bvFTD patients are sleep fragmentation and daytime sleepiness that may or may not be accompanied by insomnia (Bonakis et al., 2014; McCarter et al., 2016; Sani et al., 2019). Our findings of limited sleep fragmentation and daytime sleepiness highlight potential differences between the fly model and human disease presentation. That said, the limited yet specific deficits caused by TDP-43 OE in MBNs provide a robust measurable sleep disruption that can serve as an organism-level output for future molecular or pharmacological intervention experiments.

In addition to observed behavioral deficits, flies with TDP-43 proteinopathy show an approximately 10% reduction in median lifespan. This may be roughly comparable to human patients where FTD exerts a subtle effect on lifespan. In humans, mean age at diagnosis is 61.9 for early onset disease (< 65 years) with mean survival after diagnosis around 8 years (Kansal et al., 2016), meaning survival is reduced in comparison with the general population in most countries were clinical data are collected (e.g., 78.8 years in the United States Murphy et al., 2021). Effects on lifespan may be far more severe than a simple comparison of life expectancy would indicate, as Loi et al. (2021) report an increase in mortality risk for FTD patients when compared with age-matched controls. From the perspective of a fly model, it is somewhat surprising to see a lifespan effect given that MBNs comprise a small fraction of total neurons in the fly brain and are often thought to be largely dispensable for viability (de Belle and Heisenberg, 1994). There is evidence that lifespan is regulated at least in part by the α/β MBNs (Lien et al., 2020), a subset of MBNs included in our model, however loss of key MB proteins such as DCO (Yamazaki et al., 2007) or overexpression of vertebrate TAU in MBNs result in severe memory deficits without affecting lifespan (Mershin et al., 2004). Thus, our model provides additional evidence that specific populations of MBNs may play a role in lifespan regulation in flies, while reproducing an important, subtle characteristic of FTD.

Sequestration of mRNAs is one the mechanisms by which TDP-43 is thought to contribute to neurodegeneration and identifying candidate mRNA targets has helped better understand disease etiology (Bjork et al., 2022; Ramaswami et al., 2013). Although our model is based on TDP-43 overexpression, we note that among the mRNAs enriched with TDP-43 in MBNs we found 12 of 15 physiological targets of TDP-43 previously identified bioinformatically as potential targets of *Drosophila* TDP-43 (*i.e.*, TBPH) based on UG-richness (Langellotti et al., 2018). In future studies, it will be interesting to explore the relationship between splicing and translational targets of TDP-43. Furthermore, the overlap and specificity of some targets for MNs versus MBNs may provide insights into shared mechanisms and neuronal vulnerability in ALS and FTD, respectively.

Functional analyses of the mRNAs associated with TDP-43 identify numerous cellular pathways including Wg and Hippo that have been previously implicated in sleep regulation in flies (Harbison et al., 2017). Additionally, our findings of dopamine receptors as candidate mRNA targets is consistent with findings that loss of mesocortical dopaminergic tracts and dopamine receptors in the frontal lobes could contribute to the behavioral symptoms in FTLD (Murley and Rowe, 2018).

For functional validation we chose to focus on *dlp*, an mRNA target of TDP-43 that we previously identified in the ALS model of TDP-43 proteinopathy, and a known regulator of Wg/Wnt signaling. Indeed, Wnt signaling is dysregulated in ALS (Chen et al., 2013; Gonzalez-Fernandez et al., 2019), and mRNAs associated with Wg/Wnt pathway are enriched in FTD patient frontal cortices (Santiago et al., 2020), however, this pathway has not yet been broadly implicated in FTD pathophysiology. Using genetic interactions, we found that *dlp* OE in MBNs mitigates TDP-43 dependent working memory deficits, as evidenced by improved alternation in the Y-maze assay. Taken together, these findings support the notion that Wg/Wnt signaling is altered in ALS/FTD and modulating its activity via Dlp mitigates behavioral deficits caused by TDP-43 proteinopathy. In future studies it will be interesting to see how different mRNA targets mitigate specific phenotypic aspects of FTD and identify ALS versus FTD specific targets of TDP-43 proteinopathies.

## MATERIALS AND METHODS

### Drosophila genetics and maintenance

Flies were maintained at 25°C in 12-hour light-dark cycle with 25-30% humidity. Specific information on *Drosophila* lines used in this study and specific experiments where each line was employed can be found in the Key Resources table (Supplementary information). For aging and lifespan studies, newly eclosed virgin male and female flies were collected and maintained on standard fly cornmeal/molasses media refreshed weekly. Flies harboring *UAS TDP-43::YFP* and *UAS dlp* transgenes for co-overexpression of TDP-43 and Dlp were generated using standard genetic recombination techniques. The presence of *UAS TDP-43::YFP* was confimed by YFP expression when crossed with the pan-neuronal driver, elav GAL4 while Dlp overexpression was confirmed using RT-qPCR (*dlp OE* = 2.45 FC, *TDP-43^WT^::YFP* ; *dlp OE* =3.09 FC, *TDP-43^G298S^::YFP* ; *dlp OE* = 1.71 FC compared to *w^1118^* controls; Figure 6-supplement 1B).

### Western Blotting

1 – 3 day old flies were collected from *SS01276* crossed with 1) *w^1118^; TDP-43^WT^::YFP*, 2) *w^1118^; TDP-43^WT^::YFP; dlp OE*, 3) *w^1118^; TDP-43^WT^::YFP mCD8::RFP*, and 4) *w^1118^* in triplicate. Heads (N = 15 for each genotype) were decapitated and homogenized in 100 µl 2X Laemmli Sample Buffer (BIO-RAD 1610737) containing 5% of 2-Mercaptoethanol (Sigma-Aldrich M3148). The homogenized protein samples were boiled for 5 min in the digital Heat Block (Benchmark), and spun for 1 min. Supernatants were collected and 10 µl of protein sample was loaded in each well of precast Mini-PROTEAN TGX 4-20% gradient Gel (BIO-RAD 4561096). Following SDS-PAGE, the proteins were transferred to Nitrocellulose membrane (BIO-RAD 1620215). After transfer, the membrane was blocked in 5% nonfat milk in PBST (PBS and 0.1% TWEEN20) and incubated overnight at 4°C with primary antibodies mouse anti-GFP Living Color (Cell Signaling Technology 2955, 1:1000) to detect TDP-43 YFP and rabbit anti-Beta-Actin (Cell Signaling Technology 4967S, 1:1000) as loading control, followed by washes in TBST and incubation with secondary antibody (IRDye® 800CW, Goat anti-Mouse D21115-25, 1:10,000) and (IRDye®680RD Goat anti-Rabbit D21207-05, 1:10,000) for 1 hour at RT. The blot was imaged using a LICOR scanner (Odyssey®DLx) and protein bands intensities were quantified with LICOR “Image Studio Lite” (Figure 7-supplement 2).

### Statistical analyses

Statistical analyses were conducted in R (v. 4.1.2, R CoreTeam, 2021) and RStudio (v. 2021.09.0+351, RStudioTeam, 2022) unless specified otherwise in specific methods. The tidyverse (Wickham et al., 2019) and ggpubr (Kassambara, 2020) packages were used for summary statistics and graphics. Other analysis-specific packages are cited in the corresponding Methods section. While data for the mutant TDP-43^G298S^ model is presented as a supplement to our main findings, statistical analyses were conducted simultaneously for both TDP-43 genotypes and p-values were corrected for multiple comparisons that included both genotypes. For any given analysis when males and females did not differ statistically or sample sizes were low (*e.g.*, histological preparations), they were pooled to increase power. Where necessary, outliers were removed prior to hypothesis testing. When data did not fit the assumptions of parametric models and non-parametric analyses were used, hypothesis testing proceeded by first using a Kruskal-Wallace to test for a difference among groups, followed by pairwise comparisons using the Wilcoxon Rank Sum Test with p-values adjusted for multiple comparisons using the false discovery rate method (Benjamini and Hochberg, 1995). Specific statistical methods are described for each assay. Summary statistics for each figure can be found in the supplemental tables file.

### Mushroom body morphological analyses

Mushroom body morphology was evaluated using membrane-targeted RFP (mCD8 RFP) driven by *SS01276* (Aso, 2021). Adult brains were dissected in cold HL-3 saline (Stewart et al., 1994), fixed for 60 minutes in 4% paraformaldehyde, then rinsed in phosphate buffered saline (PBS, 3X), permeabilized in PBS with 0.25% Triton X-100 (PBST), and blocked in PBST plus 5% normal goat serum (Sigma-Aldrich 566380) 2% bovine serum albumin (Sigma-Aldrich A5611) for 45 minutes prior to antibody labeling. YFP was detected by incubating brains overnight at 4°C with a mouse monoclonal anti-GFP FITC antibody (1:300, Rockland 600-302-215). mCD8 RFP was visualized using native fluorescence. All brains were mounted on slides with the ventral side containing the mushroom body lobes (MBLs) facing the coverslip. For cell body imaging, brains were additionally incubated in Hoechst (1:10,000, Invitrogen H3570) for 10 minutes and mounted on slides with the dorsal side containing the calyx facing the coverslip.

### Image acquisition and analysis

#### TDP-43 cytoplasmic localization and MBN cell loss

Images were acquired using a Zeiss 880 Laser Scanning Confocal inverted microscope with a Plan-Apochromat 63x/1.4 oil DIC M27 lens. TDP-43 YFP signal intensity was quantified per unit area in the nucleus and cell body. Hoechst was used to define the nuclear boundaries, while total cellular TDP-43 signal was measured by tracing a boundary around YFP signal in the entire cell (Figure 1-supplement 3). To quantify the total number of MBNs expressing TDP-43 YFP, all nuclei were counted from optical sections 2 μm apart to ensure nuclei were counted only once. We tested for differences in TDP-43 nuclear to cellular ratio and MBN number by age, sex, and genotype using Analysis of Variance (ANOVA) performed on a linear model (*lm* function in R; R Core Team 2017). Residuals were normally distributed (Shapiro-Wilks test for normality, P = 0.890 and P = 0.395 for ratio and MBN number, respectively) and we therefore continued with pairwise comparisons using Tukey’s Honest Significant Difference corrected for multiple comparisons.

#### Signal intensity

All images were acquired using a Zeiss 880 Laser Scanning Confocal inverted microscope with a Plan-Apochromat 63x/1.4 oil DIC M27 lens. For mCD8 RFP fluorescence intensity, brains were imaged using either an Alexa Fluor 568 or a DS Red filter set with pinhole adjusted to 1 AU and 3 μm optical section thickness. For measuring anti-GFP FITC fluorescence intensity, brains were imaged using the FITC filter set with the pinhole adjusted to 1 AU and 0.5 μm optical section thickness. Fluorescence intensity was measured in FIJI as the integrated density over the sample area in axons by manually tracing α/β or γ lobes and subtracting adjacent background signal. First, a polygon was traced free-hand over the visible area of each lobe (α/β or γ). This polygon was then dragged to an adjacent area of the brain to obtain a mean fluorescence intensity for the background in each section. In instances where the polygon was shaped in such a way that it could not be dragged to adjacent brain area, a comparably-sized polygon was drawn in the adjacent brain region. Intensity was measured in at least three sections to obtain mean fluorescence intensity in each channel (RFP or FITC) for both α/β and γ lobes of each brain. To test for differences in signal intensity within a genotype across age time points, values of signal intensity were normalized to mean signal intensity of young (1-3 day old) brains of the same genotype and lobe. Statistical analysis of change in fluorescence intensity with age was performed by comparing middle aged (∼30 days) or old (∼60 days) flies and young (∼1 day) flies from the same genotype using the Wilcoxon Rank Sum test. Summary statistics can be found in Supplemental table S2A.

#### YFP particle size

The Analyze Particles function in FIJI was used to identify YFP puncta in MB lobes. Images were first contrast enhanced using a brightness/contrast enhancement (saturated pixels set to 0.3% and normalized) followed by thresholding using the Bernsen method in the Auto Local Threshold function with a 25 – 50 pixel radius depending on image quality. Particles were considered puncta when they were 0.25 – 3.0 μm^2^ in size and showed 0.25 - 1.0 circularity. To measure mean particle size in each sample, a polygon was drawn over the visible portion of each lobe in at least three sections for each lobe. From these subsamples we calculated a mean particle size for each lobe of each brain. Summary statistics can be found in Supplemental table S2B.

#### GAL4 dilution effect

To test whether UAS-driven expression of a second transgene reduces UAS-driven TDP-43^WT^ expression, in addition to western blots, we also measured mean pixel intensity from maximum intensity projections of the horizontal lobes (β and γ combined in maximum intensity projection) in two genotypes with *SS01276*-driven transgenes, *OR-R ;TDP-43^WT^::YFP* and *w^1118^ ; TDP-43^WT^::YFP mCD8::RFP* (Figure 7-supplement 3).

### Behavioral assays

#### Y-maze

To measure changes in working memory, we employed a y-maze assay described by Lewis et al. (2017). We tracked the two-dimensional movement of individual *Drosophila* placed in small symmetrical Y-mazes using video recordings for ten 1-minute trials following two minutes of acclimation time in the maze. Observations took place between 08:45 and 11:00 and 14:00 to 16:00 when flies were most active. Male and female flies were run separately and the locations of genotypes in the 7 by 7 maze array were randomized for each set of trials. Assays were conducted in a dark room with each maze lit uniformly from below with a white LED array and capped with a clear Plexiglas coverslip. The overall movement of flies and visits to three unique arms consecutively were quantified using the Noldus Ethovision software. Spontaneous alternation behavior was measured by scoring three consecutive arm entries in a sliding window for each of ten trials. The alternation score was calculated as alternations divided by alternation attempts. Movement and alternation data were summed for each fly during each 1-minute trial and then averaged over ten trials for statistical analyses. Flies originating from three biological replicates were pooled for analysis (Supplementary table S3). The large samples sizes permitted the use of parametric tests, and we assessed differences in distance moved or alternation score across genotypes using ANOVA on a linear mixed-effects model (lmer function in R; Bates et al., 2015) that included replicate as a fixed effect. Flies that performed no alternations naturally did not have a percent alternation score and were removed from subsequent analysis. Males and females were analyzed separately. Flies were additionally assessed for the presence of bias for specific arms, calculated as a ratio of the mean number of entries into each arm over mean total entries and tested for significant variation from 0.333 using the Wilcoxon Signed Rank test. Interestingly, while females showed no arm biases, males overexpressing *TDP-43^WT^* and OR-R controls showed a bias against arm B (*TDP-43^WT^*, 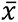 = 0.307 ± 0.08, P < 0.0001; OR-R, 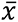 = 0.316 ± 0.13, P = 0.0038). *TDP-43^WT^* also showed a bias for arm A (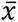 = 0.352 ± 0.076, P = 0.0033), while OR-R controls showed a bias for arm C (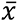] = 0.347 ± 0.113, P = 0.048). Although these analyses suggest that arm biases exist in males and are not unique to TDP-43 overexpression, a single replicate was used to test whether Dlp overexpression could rescue alternation deficits seen in TDP-43 overexpressing flies therefore hypothesis testing proceeded with non-parametric tests as described in *Statistical analyses*.

#### Sleep

Adult flies from the three age time points were monitored individually using *Drosophila* Activity Monitors (DAMs; Trikinetics, Waltham, MA; Chiu et al., 2010). Flies were placed in monitoring tubes with a small amount of food the day before they reached 2-3, 31-33, or 61-63 days old. Sleep and activity monitoring began at 12 AM following placement of flies in the sleep incubator, allowing flies at least six hours prior to the start of the experiment to acclimate to the tubes. Sleep and locomotor activity data were collected at 1-minute intervals for three days and analyzed using ShinyR-DAM (Cichewicz and Hirsh, 2018). In *Drosophila*, sleep is defined at 5 minutes of inactivity (Shaw et al., 2000) and here sleep data were processed using the ShinyR DAM program (Cichewicz and Hirsh 2018), which provided average sleep, sleep bout number and duration, and activity to sleep bout length ratio. Comparison of these variables by age and genotype were conducted separately for male and female flies. Data from two replicates were pooled (> 15 flies per genotype x replicate; sample sizes in supplemental tables S4A, S4B).

### Lifespan

Newly eclosed virgin males and females were separated and placed on fresh food, then transferred weekly into new food vials. The number of surviving flies was counted every other day for 100 days. Survival analysis and plots were generated using R packages *survival* (Therneau, 2021) and *survivminer* (Kassambara et al., 2021).

### mRNA targets of TDP-43

#### RNA Immunoprecipitations

Equal numbers of male and female flies, aged 1-3 days, were pooled in Eppendorf tubes and flash frozen in liquid nitrogen. A minimum of 500 flies expressing YFP, TDP-43^WT^ YFP, or TDP-43^G298S^ YFP were collected for each of three biological replicates. Frozen flies were then transferred to 50mL conical tubes and heads were separated from the bodies using four rounds of vortexing and flash freezing with liquid nitrogen. A sieve was used to filter out the bodies and isolated heads were collected into tubes containing lysis beads (Next Advance Green lysis beads). Heads were homogenized (Next Advance Bullet Blender) in 1mL fresh lysis buffer (DEPC water, 50 mM HEPES buffer pH 7.4, 0.5% Triton X-100, 150 mM NaCl, 30 mM EDTA), protease inhibitors (cOmplete™, Mini, EDTA-free Protease Inhibitor Cocktail, Millipore Sigma 11873580001), and RNAsin Plus 400 u/mL (Fischer Scientific PRN2615), centrifuged for 10 minutes at 10,000 rpm, and the lysate was collected. A portion of the lysate was saved for the protein input and RNA input samples. For protein input, the lysate was mixed with 2X Laemmli buffer, boiled at 95-100°C for 10 minutes, and stored at -20°C. For RNA input, TRIzol reagent (Thermofisher 15596026) was added to the lysate and the sample was stored at -80°C. High affinity GFP-Trap magnetic agarose beads (ChromoTek) were added to the remaining lysate and rotated end-over-end for 90 minutes at 4°C to allow for binding of YFP. The beads were separated from the lysate with a magnet and washed three times with fresh wash buffer (DEPC water, 50 mM HEPES buffer pH 7.4, 0.5% Triton X-100, 150 mM NaCl, 30 mM EDTA). The beads were resuspended in wash buffer and split into two tubes for the protein IP and RNA IP samples. For the protein IP sample, 2X Laemmli buffer was added to the beads, samples were boiled at 95-100°C for 10 minutes, and the beads were removed with a magnet. Western blots (see above) were performed to ensure that the immunoprecipitated complexes contained TDP-43 before processing the RNA IPs. For the RNA IP sample, TRIzol reagent was added to the beads, the solution was pipetted up and down for 60 seconds, and the beads were removed with a magnet.

#### RNA-Seq

RNA was quantified by nanodrop to bring it within range for ribogreen quantification. RNA was also checked using Agilent Tapestation High Sensitivitiy RNA screentape. 1 ng total RNA was used for SMART-Seq HT PLUS (Takara Bio USA, Inc. Cat # R400748) following manufacturer’s protocol. Determination of cDNA quality and quantity was determined via Agilent Tapestation High Sensitivity D5000 Screentape and Qubit dsDNA HS assay for input into library amplification. Libraries were quantified by Agilent Tapestation High Sensitivity D1000 Screentape and Kapa Library Quantification kit for Illumina platforms. Libraries were pooled equimolarly, pools were quantified by Agilent Tapestation High Sensitivity D1000 Screentape and Kapa Library Quantification kit and loaded on the NovaSeq 6000 S4 flowcell and sequenced to 101 x 11 x 11 x 101 cycles. Trimmed fastqs were aligned to the Dmel genome with STAR v2.6.1d (Dobin et al., 2013). Aligned reads were counted with featureCounts v1.6.3 (Liao et al., 2014) using the genome annotation. Files produced for individual samples were aggregated into a single .txt file using python. Differential expression was quantified using DESeq2 (Love et al., 2014). We included overexpression of cytoplasmic YFP in MBNs as a control for IPs, however the expression levels of YFP alone were far greater than cytoplasmic TDP-43 YFP (data not shown), making it difficult to assess the role of cytoplasmic YFP alone. To ensure that candidate mRNA targets showing the greatest enrichment were indeed specific in their association with TDP-43, we subtracted YFP control Log2FC values from the Log2FC values of targets recovered in each of our models. Indeed, in this YFP-subtracted analysis *futsch* was recovered as enriched with TDP-43 in both variants (Figure 6A, Figure 6-figure supplement 1A) and the majority of mRNAs that showed high enrichment in the original analysis were retained in the YFP-subtracted analyses (Figure 6A vs 6D, Supplemental figure S6A vs S6C).

#### Functional annotation

The Database for Annotation, Visualization, and Integrated Discovery (DAVID) was used to functionally annotated genes enriched with TDP-43 (Dennis et al., 2003; Sherman et al., 2022). To assess genes over-represented in our disease model data sets, we used lists of genes significantly enriched with each of our TDP-43 models (differential gene expression lists available in source data).

#### RNA Extraction for Validation

RNA input and RNA IP samples, stored in TRIzol reagent (Thermofisher 15596026) at -80°C, were allowed to thaw completely on ice and chloroform was added to each sample. Samples were shaken briefly to mix, left to incubate for 3 minutes at room temperature, and then centrifuged at 12,000 rpm for 15 minutes at 4°C to allow for separation of the aqueous and organic phases. The RNA-containing aqueous phase was collected and molecular grade isopropyl alcohol was added to each sample. Samples were incubated for 10 minutes at room temperature for RNA precipitation and then centrifuged at 12,000 rpm for 10 minutes. The pelleted RNA was washed with 75% ethanol (200 proof ethanol & HyPure water) and left to dry in a fume hood. RNA was resuspended in HyPure water and the concentration and 260/280 absorbance ratio were measured using Nanodrop.

#### RT-qPCR

RNA extracted from input and IP samples was used as template RNA for cDNA synthesis by reverse transcription. Total RNA used in the cDNA synthesis reactions was normalized across samples. cDNA was synthesized using the Fisher First Strand cDNA synthesis kit (Thermofisher Scientific K1641). qPCR reactions were prepared in a 96-well qPCR plate, in triplicates, using Taqman Fast Advanced Master Mix (Thermofisher Scientific 4444556) and Taqman probes for Dally-like protein (dlp) (Thermofisher Scientific Dm01798597_m1) and Gpdh1 (Thermofisher Scientific Dm01841185_m1). qPCR was conducted on the qTOWER (Analytik Jena 844-00504-4) qPCR machine. Delta Ct was calculated as the difference in Ct values (dlp - gpdh1). To determine enrichment of dlp mRNA in the IPs, ΔΔCt was calculated as the difference in Delta Ct values (IP - Input). Fold change was calculated as 2^(-ΔΔCt).

#### Dally-like protein target validation

Brains from flies 1-3 days or 50-55 days were dissected out and the tissue fixed, permeabilized and blocked as described above. Brains were then incubated overnight at 4°C in an anti-Dally-like protein antibody at 1:5 in block (13G8 developed by Phil Beachy, obtained from the Developmental Studies Hybridoma Bank, created by the NICHD of the NIH and maintained at The University of Iowa, Department of Biology, Iowa City, IA 52242, targets amino acids V523 to Q702), rinsed 3X in 0.1% PBST and incubated in goat anti-mouse Alexa Flour 647 antibody overnight at 4°C (1:300, ThermoFisher A32728). Images were acquired using a Zeiss 880 Laser Scanning Confocal inverted microscope with a Plan-Apochromat 63x/1.4 oil DIC M27 lens. Signal intensity was traced from maximum intensity projections of the MBLs and normalized to background intensity. Dlp signal intensity declined with age in TDP-43 flies, so the mCD8-RFP channel was used to trace the lobes in aged brains, ensuring the correct lobe area was used for signal intensity measurements.

## DATA AVAILABILITY

Summary statistics for each figure can be found in the supplemental tables file. Source data are available from ScholarSphere (doi:10.26207/jq6p-w169) and are cataloged in the Key Resources table. Please refer to the included readme file for variable descriptions. RNA-seq data are available from NCBI GEO Bioproject GSE217213.

## ACKNOWLEDGEMENTS

We would like to thank Patty Jansma at the University of Arizona Imaging Cores - Optical Facility for her consultation on sample preparation and image acquisition and Dr. Shaun Davis in the Schlenke lab at UArizona for generating TDP-43 transgenic lines in a OR-R background. We thank Erik Lehmkuhl for his advice on immunoprecipitation experiments and interpretation of RNA-seq data. We also acknowledge Dr. Robert Kraft for generating UAS TDP-43 UAS mCD8 RFP recombinant stocks. We thank Dr. Yoshi Aso (HHMI) for sharing the *SS01276* GAL4 line and Dr. Fabian Fernandez (UArizona) for Y-maze training. Support for Hillary Cowell Ruvalcaba was provided in part by The University of Arizona’s Undergraduate Research Opportunities Consortium Prep (UROC-Prep) Program. The University of Arizona Maximizing Access to Research Careers (MARC Training Grant NIGMS T34 GM008718) program provided support for Grace Hala’ufia. We would like to acknowledge members of the Zarnescu lab for their advice on experimental design and data analysis, and feedback on the manuscript.

## REFERENCES

Al-Chalabi, A., Jones, A., Troakes, C., King, A., Al-Sarraj, S., and van den Berg, L.H. (2012). The genetics and neuropathology of amyotrophic lateral sclerosis. Acta Neuropathol 124, 339–352. 10.1007/s00401-012-1022-4.

Alami, N.H., Smith, R.B., Carrasco, M.A., Williams, L.A., Winborn, C.S., Han, S.S., Kiskinis, E., Winborn, B., Freibaum, B.D., Kanagaraj, A., et al. (2014). Axonal transport of TDP-43 mRNA granules is impaired by ALS-causing mutations. Neuron 81, 536–543. 10.1016/j.neuron.2013.12.018.

Altman, T., Ionescu, A., Ibraheem, A., Priesmann, D., Gradus-Pery, T., Farberov, L., Alexandra, G., Shelestovich, N., Dafinca, R., Shomron, N., et al. (2021). Axonal TDP-43 condensates drive neuromuscular junction disruption through inhibition of local synthesis of nuclear encoded mitochondrial proteins. Nat Commun 12, 6914. 10.1038/s41467-021-27221-8.

Amador-Ortiz, C., Lin, W.L., Ahmed, Z., Personett, D., Davies, P., Duara, R., Graff-Radford, N.R., Hutton, M.L., and Dickson, D.W. (2007). TDP-43 immunoreactivity in hippocampal sclerosis and Alzheimer’s disease. Ann Neurol 61, 435–445. 10.1002/ana.21154.

Arai, T., Hasegawa, M., Akiyama, H., Ikeda, K., Nonaka, T., Mori, H., Mann, D., Tsuchiya, K., Yoshida, M., Hashizume, Y., and Oda, T. (2006). TDP-43 is a component of ubiquitin-positive tau-negative inclusions in frontotemporal lobar degeneration and amyotrophic lateral sclerosis. Biochem Biophys Res Commun 351, 602–611. 10.1016/j.bbrc.2006.10.093.

Ash, P.E., Zhang, Y.J., Roberts, C.M., Saldi, T., Hutter, H., Buratti, E., Petrucelli, L., and Link, C.D. (2010). Neurotoxic effects of TDP-43 overexpression in C. elegans. Hum Mol Genet 19, 3206–3218. 10.1093/hmg/ddq230.

Aso, Y. (2021). Split-GAL4 Mushroom Body Driver Line

Aso, Y., Grubel, K., Busch, S., Friedrich, A.B., Siwanowicz, I., and Tanimoto, H. (2009). The mushroom body of adult Drosophila characterized by GAL4 drivers. J Neurogenet 23, 156–172. 10.1080/01677060802471718.

Aso, Y., Hattori, D., Yu, Y., Johnston, R.M., Iyer, N.A., Ngo, T.T., Dionne, H., Abbott, L.F., Axel, R., Tanimoto, H., and Rubin, G.M. (2014). The neuronal architecture of the mushroom body provides a logic for associative learning. Elife 3, e04577. 10.7554/eLife.04577.

Ayala, Y.M., Pantano, S., D’Ambrogio, A., Buratti, E., Brindisi, A., Marchetti, C., Romano, M., and Baralle, F.E. (2005). Human, Drosophila, and C.elegans TDP43: nucleic acid binding properties and splicing regulatory function. J Mol Biol 348, 575-588. 10.1016/j.jmb.2005.02.038.

Azpurua, J., El-Karim, E.G., Tranquille, M., and Dubnau, J. (2021). A behavioral screen for mediators of age-dependent TDP-43 neurodegeneration identifies SF2/SRSF1 among a group of potent suppressors in both neurons and glia. PLoS Genet 17, e1009882. 10.1371/journal.pgen.1009882.

Baier, A., Wittek, B., and Brembs, B. (2002). Drosophila as a new model organism for the neurobiology of aggression? J Exp Biol 205, 1233–1240. 10.1242/jeb.205.9.1233.

Bates, D., Mächler, M., Bolker, B., and Walker, S. (2015). Fitting Linear Mixed-Effects Models Usinglme4. Journal of Statistical Software 67. 10.18637/jss.v067.i01.

Bellen, H.J. (1998). The fruit fly: a model organism to study the genetics of alcohol abuse and addiction? Cell 93, 909–912. 10.1016/s0092-8674(00)81195-2.

Benjamini, Y., and Hochberg, Y. (1995). Controlling the False Discovery Rate A Practical and Powerful Approach to Multiple Testing. Journal of the Royal Statistical Society. Series B (Methodological) 57, 289–300.

Bier, E. (2005). Drosophila, the golden bug, emerges as a tool for human genetics. Nat Rev Genet 6, 9–23. 10.1038/nrg1503.

Bjork, R.T., Mortimore, N.P., Loganathan, S., and Zarnescu, D.C. (2022). Dysregulation of Translation in TDP-43 Proteinopathies: Deficits in the RNA Supply Chain and Local Protein Production. Front Neurosci 16, 840357. 10.3389/fnins.2022.840357.

Bonakis, A., Economou, N.T., Paparrigopoulos, T., Bonanni, E., Maestri, M., Carnicelli, L., Di Coscio, E., Ktonas, P., Vagiakis, E., Theodoropoulos, P., and Papageorgiou, S.G. (2014). Sleep in frontotemporal dementia is equally or possibly more disrupted, and at an earlier stage, when compared to sleep in Alzheimer’s disease. J Alzheimers Dis 38, 85–91. 10.3233/JAD-122014.

Borroni, B., Bonvicini, C., Alberici, A., Buratti, E., Agosti, C., Archetti, S., Papetti, A., Stuani, C., Di Luca, M., Gennarelli, M., and Padovani, A. (2009). Mutation within TARDBP leads to Frontotemporal Dementia without motor neuron disease. Human Mutation 30, E974–E983. 10.1002/humu.21100.

Bridi, J.C., Ludlow, Z.N., Kottler, B., Hartmann, B., Vanden Broeck, L., Dearlove, J., Goker, M., Strausfeld, N.J., Callaerts, P., and Hirth, F. (2020). Ancestral regulatory mechanisms specify conserved midbrain circuitry in arthropods and vertebrates. Proc Natl Acad Sci U S A 117, 19544–19555. 10.1073/pnas.1918797117.

Cairns, N.J., Neumann, M., Bigio, E.H., Holm, I.E., Troost, D., Hatanpaa, K.J., Foong, C., White, C.L., 3rd, Schneider, J.A., Kretzschmar, H.A., et al. (2007). TDP-43 in familial and sporadic frontotemporal lobar degeneration with ubiquitin inclusions. Am J Pathol 171, 227-240. 10.2353/ajpath.2007.070182.

Chakraborty, R., Vepuri, V., Mhatre, S.D., Paddock, B.E., Miller, S., Michelson, S.J., Delvadia, R., Desai, A., Vinokur, M., Melicharek, D.J., et al. (2011). Characterization of a Drosophila Alzheimer’s disease model: pharmacological rescue of cognitive defects. PLoS One 6, e20799. 10.1371/journal.pone.0020799.

Chang, X.-L., Tan, M.-S., Tan, L., and Yu, J.-T. (2015). The Role of TDP-43 in Alzheimer’s Disease. Molecular Neurobiology 53, 3349–3359. 10.1007/s12035-015-9264-5.

Chen, H.J., Topp, S.D., Hui, H.S., Zacco, E., Katarya, M., McLoughlin, C., King, A., Smith, B.N., Troakes, C., Pastore, A., and Shaw, C.E. (2019). RRM adjacent TARDBP mutations disrupt RNA binding and enhance TDP-43 proteinopathy. Brain 142, 3753–3770. 10.1093/brain/awz313.

Chen, Y., Guan, Y., Zhang, Z., Liu, H., Wang, S., Yu, L., Wu, X., and Wang, X. (2013). Wnt signaling pathway is involved in the pathogenesis of amyotrophic lateral sclerosis in adult transgenic mice. Neurological Research 34, 390–399. 10.1179/1743132812y.0000000027.

Chiu, J.C., Low, K.H., Pike, D.H., Yildirim, E., and Edery, I. (2010). Assaying locomotor activity to study circadian rhythms and sleep parameters in Drosophila. J Vis Exp. 10.3791/2157.

Chou, C.-C., Zhang, Y., Umoh, M.E., Vaughan, S.W., Lorenzini, I., Liu, F., Sayegh, M., Donlin-Asp, P.G., Chen, Y.H., Duong, D.M., et al. (2018). TDP-43 pathology disrupts nuclear pore complexes and nucleocytoplasmic transport in ALS/FTD. Nature Neuroscience 21, 228–239. 10.1038/s41593-017-0047-3.

Cichewicz, K., and Hirsh, J. (2018). ShinyR-DAM: a program analyzing Drosophila activity, sleep and circadian rhythms. Commun Biol 1, 25. 10.1038/s42003-018-0031-9.

Colombrita, C., Zennaro, E., Fallini, C., Weber, M., Sommacal, A., Buratti, E., Silani, V., and Ratti, A. (2009). TDP-43 is recruited to stress granules in conditions of oxidative insult. J Neurochem 111, 1051–1061. 10.1111/j.1471-4159.2009.06383.x.

Coyne, A.N., Lorenzini, I., Chou, C.C., Torvund, M., Rogers, R.S., Starr, A., Zaepfel, B.L., Levy, J., Johannesmeyer, J., Schwartz, J.C., et al. (2017). Post-transcriptional Inhibition of Hsc70-4/HSPA8 Expression Leads to Synaptic Vesicle Cycling Defects in Multiple Models of ALS. Cell Rep 21, 110–125. 10.1016/j.celrep.2017.09.028.

Coyne, A.N., Siddegowda, B.B., Estes, P.S., Johannesmeyer, J., Kovalik, T., Daniel, S.G., Pearson, A., Bowser, R., and Zarnescu, D.C. (2014). Futsch/MAP1B mRNA is a translational target of TDP-43 and is neuroprotective in a Drosophila model of amyotrophic lateral sclerosis. J Neurosci 34, 15962–15974. 10.1523/JNEUROSCI.2526-14.2014.

Coyne, A.N., Yamada, S.B., Siddegowda, B.B., Estes, P.S., Zaepfel, B.L., Johannesmeyer, J.S., Lockwood, D.B., Pham, L.T., Hart, M.P., Cassel, J.A., et al. (2015). Fragile X protein mitigates TDP-43 toxicity by remodeling RNA granules and restoring translation. Hum Mol Genet 24, 6886–6898. 10.1093/hmg/ddv389.

Crittenden (1998). Tripartite Mushroom Body Architecture Revealed by Antigenic Markers.

de Belle, J.S., and Heisenberg, M. (1994). Associative odor learning in Drosophila abolished by chemical ablation of mushroom bodies. Science 263, 692–695. 10.1126/science.8303280.

Dennis, G., Jr., Sherman, B.T., Hosack, D.A., Yang, J., Gao, W., Lane, H.C., and Lempicki, R.A. (2003). DAVID: Database for Annotation, Visualization, and Integrated Discovery. Genome Biol 4, P3.

Dobin, A., Davis, C.A., Schlesinger, F., Drenkow, J., Zaleski, C., Jha, S., Batut, P., Chaisson, M., and Gingeras, T.R. (2013). STAR: ultrafast universal RNA-seq aligner. Bioinformatics 29, 15–21. 10.1093/bioinformatics/bts635.

Driscoll, M., Buchert, S.N., Coleman, V., McLaughlin, M., Nguyen, A., and Sitaraman, D. (2021). Compartment specific regulation of sleep by mushroom body requires GABA and dopaminergic signaling. Sci Rep 11, 20067. 10.1038/s41598-021-99531-2.

Elden, A.C., Kim, H.J., Hart, M.P., Chen-Plotkin, A.S., Johnson, B.S., Fang, X., Armakola, M., Geser, F., Greene, R., Lu, M.M., et al. (2010). Ataxin-2 intermediate-length polyglutamine expansions are associated with increased risk for ALS. Nature 466, 1069–1075. 10.1038/nature09320.

Enell, L.E., Kapan, N., Soderberg, J.A., Kahsai, L., and Nassel, D.R. (2010). Insulin signaling, lifespan and stress resistance are modulated by metabotropic GABA receptors on insulin producing cells in the brain of Drosophila. PLoS One 5, e15780. 10.1371/journal.pone.0015780.

Erwin, D.H. (2015). Early metazoan life: divergence, environment and ecology. Philos Trans R Soc Lond B Biol Sci 370. 10.1098/rstb.2015.0036.

Estes, P.S., Boehringer, A., Zwick, R., Tang, J.E., Grigsby, B., and Zarnescu, D.C. (2011). Wild-type and A315T mutant TDP-43 exert differential neurotoxicity in a Drosophila model of ALS. Hum Mol Genet 20, 2308–2321. 10.1093/hmg/ddr124.

Estes, P.S., Daniel, S.G., McCallum, A.P., Boehringer, A.V., Sukhina, A.S., Zwick, R.A., and Zarnescu, D.C. (2013). Motor neurons and glia exhibit specific individualized responses to TDP-43 expression in a Drosophila model of amyotrophic lateral sclerosis. Dis Model Mech 6, 721–733. 10.1242/dmm.010710.

Fallini, C., Bassell, G.J., and Rossoll, W. (2012). The ALS disease protein TDP-43 is actively transported in motor neuron axons and regulates axon outgrowth. Hum Mol Genet 21, 3703–3718. 10.1093/hmg/dds205.

Feiguin, F., Godena, V.K., Romano, G., D’Ambrogio, A., Klima, R., and Baralle, F.E. (2009). Depletion of TDP-43 affects Drosophila motoneurons terminal synapsis and locomotive behavior. FEBS Lett 583, 1586–1592. 10.1016/j.febslet.2009.04.019.

Ferrari, R., Kapogiannis, D., Huey, E.D., and Momeni, P. (2011). FTD and ALS: a tale of two diseases. Curr Alzheimer Res 8, 273–294. 10.2174/156720511795563700.

Fiesel, F.C., Weber, S.S., Supper, J., Zell, A., and Kahle, P.J. (2012). TDP-43 regulates global translational yield by splicing of exon junction complex component SKAR. Nucleic Acids Res 40, 2668–2682. 10.1093/nar/gkr1082.

Freibaum, B.D., Chitta, R.K., High, A.A., and Taylor, J.P. (2010). Global analysis of TDP-43 interacting proteins reveals strong association with RNA splicing and translation machinery. J Proteome Res 9, 1104–1120. 10.1021/pr901076y.

Gao, J., Wang, L., Huntley, M.L., Perry, G., and Wang, X. (2018). Pathomechanisms of TDP-43 in neurodegeneration. J Neurochem. 10.1111/jnc.14327.

Gonzalez-Fernandez, C., Gonzalez, P., Andres-Benito, P., Ferrer, I., and Rodriguez, F.J. (2019). Wnt Signaling Alterations in the Human Spinal Cord of Amyotrophic Lateral Sclerosis Cases: Spotlight on Fz2 and Wnt5a. Mol Neurobiol 56, 6777–6791. 10.1007/s12035-019-1547-9.

Graham, A., Davies, R., Xuereb, J., Halliday, G., Kril, J., Creasey, H., Graham, K., and Hodges, J. (2005). Pathologically proven frontotemporal dementia presenting with severe amnesia. Brain 128, 597–605. 10.1093/brain/awh348.

Hampel, S., Chung, P., McKellar, C.E., Hall, D., Looger, L.L., and Simpson, J.H. (2011). Drosophila Brainbow: a recombinase-based fluorescence labeling technique to subdivide neural expression patterns. Nat Methods 8, 253–259. 10.1038/nmeth.1566.

Harbison, S.T., Serrano Negron, Y.L., Hansen, N.F., and Lobell, A.S. (2017). Selection for long and short sleep duration in Drosophila melanogaster reveals the complex genetic network underlying natural variation in sleep. PLoS Genet 13, e1007098. 10.1371/journal.pgen.1007098.

Hasegawa, M., Arai, T., Akiyama, H., Nonaka, T., Mori, H., Hashimoto, T., Yamazaki, M., and Oyanagi, K. (2007). TDP-43 is deposited in the Guam parkinsonism-dementia complex brains. Brain 130, 1386–1394. 10.1093/brain/awm065.

Heisenberg, M. (2003). Mushroom body memoir: from maps to models. Nat Rev Neurosci 4, 266–275. 10.1038/nrn1074.

Hornberger, M., and Piguet, O. (2012). Episodic memory in frontotemporal dementia: a critical review. Brain 135, 678–692. 10.1093/brain/aws011.

Isabel, G., Pascual, A., and Preat, T. (2004). Exclusive consolidated memory phases in Drosophila. Science 304, 1024–1027. 10.1126/science.1094932.

Ito, K., Awano, W., Suzuki, K., Hiromi, Y., and Yamamoto, D. (1997). The Drosophila mushroom body is a quadruple structure of clonal units each of which contains a virtually identical set of neurones and glial cells. Development 124, 761–771. 10.1242/dev.124.4.761.

Jo, M., Lee, S., Jeon, Y.M., Kim, S., Kwon, Y., and Kim, H.J. (2020). The role of TDP-43 propagation in neurodegenerative diseases: integrating insights from clinical and experimental studies. Exp Mol Med 52, 1652–1662. 10.1038/s12276-020-00513-7.

Joiner, W.J., Crocker, A., White, B.H., and Sehgal, A. (2006). Sleep in Drosophila is regulated by adult mushroom bodies. Nature 441, 757–760. 10.1038/nature04811.

Josephs, K.A., Murray, M.E., Whitwell, J.L., Parisi, J.E., Petrucelli, L., Jack, C.R., Petersen, R.C., and Dickson, D.W. (2014a). Staging TDP-43 pathology in Alzheimer’s disease. Acta Neuropathol 127, 441–450. 10.1007/s00401-013-1211-9.

Josephs, K.A., Whitwell, J.L., Weigand, S.D., Murray, M.E., Tosakulwong, N., Liesinger, A.M., Petrucelli, L., Senjem, M.L., Knopman, D.S., Boeve, B.F., et al. (2014b). TDP-43 is a key player in the clinical features associated with Alzheimer’s disease. Acta Neuropathol 127, 811–824. 10.1007/s00401-014-1269-z.

Kabashi, E., Valdmanis, P.N., Dion, P., Spiegelman, D., McConkey, B.J., Vande Velde, C., Bouchard, J.P., Lacomblez, L., Pochigaeva, K., Salachas, F., et al. (2008). TARDBP mutations in individuals with sporadic and familial amyotrophic lateral sclerosis. Nat Genet 40, 572–574. 10.1038/ng.132.

Kamyshev, N.G., Smirnova, G.P., Kamysheva, E.A., Nikiforov, O.N., Parafenyuk, I.V., and Ponomarenko, V.V. (2002). Plasticity of social behavior in Drosophila. Neurosci Behav Physiol 32, 401–408. 10.1023/a:1015832328023.

Kansal, K., Mareddy, M., Sloane, K.L., Minc, A.A., Rabins, P.V., McGready, J.B., and Onyike, C.U. (2016). Survival in Frontotemporal Dementia Phenotypes: A Meta-Analysis. Dement Geriatr Cogn Disord 41, 109–122. 10.1159/000443205.

Kassambara, A. (2020). ggpubr: ’ggplot2’ Based Publication Ready Plots.

Kassambara, A., Kosinski, M., and Biecek, P. (2021). survminer: Drawing Survival Curves using ’ggplot2’. R package version 0.4.9.

Khalfallah, Y., Kuta, R., Grasmuck, C., Prat, A., Durham, H.D., and Vande Velde, C. (2018). TDP-43 regulation of stress granule dynamics in neurodegenerative disease-relevant cell types. Sci Rep 8, 7551. 10.1038/s41598-018-25767-0.

Kunz, T., Kraft, K.F., Technau, G.M., and Urbach, R. (2012). Origin of Drosophila mushroom body neuroblasts and generation of divergent embryonic lineages. Development 139, 2510–2522. 10.1242/dev.077883.

Langellotti, S., Romano, G., Feiguin, F., Baralle, F.E., and Romano, M. (2018). RhoGAPp190: A potential player in tbph-mediated neurodegeneration in Drosophila. PLoS One 13, e0195845. 10.1371/journal.pone.0195845.

Lee, T., Lee, A., and Luo, L. (1999). Development of the Drosophila mushroom bodies: sequential generation of three distinct types of neurons from a neuroblast. Development 126, 4065–4076. 10.1242/dev.126.18.4065.

Lehmkuhl, E.M., Loganathan, S., Alsop, E., Blythe, A.D., Kovalik, T., Mortimore, N.P., Barrameda, D., Kueth, C., Eck, R.J., Siddegowda, B.B., et al. (2021). TDP-43 proteinopathy alters the ribosome association of multiple mRNAs including the glypican Dally-like protein (Dlp)/GPC6. Acta Neuropathol Commun 9, 52. 10.1186/s40478-021-01148-z.

Lewis, S.A., Negelspach, D.C., Kaladchibachi, S., Cowen, S.L., and Fernandez, F. (2017). Spontaneous alternation: A potential gateway to spatial working memory in Drosophila. Neurobiol Learn Mem 142, 230–235. 10.1016/j.nlm.2017.05.013.

Li, J., Vitiello, M.V., and Gooneratne, N.S. (2018). Sleep in Normal Aging. Sleep Med Clin 13, 1–11. 10.1016/j.jsmc.2017.09.001.

Li, Y., Ray, P., Rao, E.J., Shi, C., Guo, W., Chen, X., Woodruff, E.A., 3rd, Fushimi, K., and Wu, J.Y. (2010). A Drosophila model for TDP-43 proteinopathy. Proc Natl Acad Sci U S A 107, 3169-3174. 10.1073/pnas.0913602107.

Liao, Y., Smyth, G.K., and Shi, W. (2014). featureCounts: an efficient general purpose program for assigning sequence reads to genomic features. Bioinformatics 30, 923–930. 10.1093/bioinformatics/btt656.

Lien, W.Y., Chen, Y.T., Li, Y.J., Wu, J.K., Huang, K.L., Lin, J.R., Lin, S.C., Hou, C.C., Wang, H.D., Wu, C.L., et al. (2020). Lifespan regulation in alpha/beta posterior neurons of the fly mushroom bodies by Rab27. Aging Cell 19, e13179. 10.1111/acel.13179.

Ling, J.P., Pletnikova, O., Troncoso, J.C., and Wong, P.C. (2015). TDP-43 repression of nonconserved cryptic exons is compromised in ALS-FTD. Science 349, 650–655. 10.1126/science.aab0983.

Ling, S.C., Polymenidou, M., and Cleveland, D.W. (2013). Converging mechanisms in ALS and FTD: disrupted RNA and protein homeostasis. Neuron 79, 416–438. 10.1016/j.neuron.2013.07.033.

Liu, L., Wolf, R., Ernst, R., and Heisenberg, M. (1999). Context generalization in Drosophila visual learning requires the mushroom bodies. Nature 400, 753–756. 10.1038/23456.

Loganathan, S., Wilson, B.A., Carey, S.B., Manzo, E., Joardar, A., Ugur, B., and Zarnescu, D.C. (2022). TDP-43 Proteinopathy Causes Broad Metabolic Alterations including TCA Cycle Intermediates and Dopamine Levels in Drosophila Models of ALS. Metabolites 12. 10.3390/metabo12020101.

Loi, S.M., Tsoukra, P., Chen, Z., Wibawa, P., Eratne, D., Kelso, W., Walterfang, M., and Velakoulis, D. (2021). Risk factors to mortality and causes of death in frontotemporal dementia: An Australian perspective. Int J Geriatr Psychiatry 37. 10.1002/gps.5668.

Love, M.I., Huber, W., and Anders, S. (2014). Moderated estimation of fold change and dispersion for RNA-seq data with DESeq2. Genome Biol 15, 550. 10.1186/s13059-014-0550-8.

Lu, Y., Ferris, J., and Gao, F.B. (2009). Frontotemporal dementia and amyotrophic lateral sclerosis-associated disease protein TDP-43 promotes dendritic branching. Mol Brain 2, 30. 10.1186/1756-6606-2-30.

Mackenzie, I.R., Bigio, E.H., Ince, P.G., Geser, F., Neumann, M., Cairns, N.J., Kwong, L.K., Forman, M.S., Ravits, J., Stewart, H., et al. (2007). Pathological TDP-43 distinguishes sporadic amyotrophic lateral sclerosis from amyotrophic lateral sclerosis with SOD1 mutations. Ann Neurol 61, 427–434. 10.1002/ana.21147.

Mackenzie, I.R., Neumann, M., Baborie, A., Sampathu, D.M., Du Plessis, D., Jaros, E., Perry, R.H., Trojanowski, J.Q., Mann, D.M., and Lee, V.M. (2011). A harmonized classification system for FTLD-TDP pathology. Acta Neuropathol 122, 111–113. 10.1007/s00401-011-0845-8.

Mackenzie, I.R.A., Rademakers, R., and Neumann, M. (2010). TDP-43 and FUS in amyotrophic lateral sclerosis and frontotemporal dementia. The Lancet Neurology 9, 995–1007. 10.1016/s1474-4422(10)70195-2.

Manzo, E., Lorenzini, I., Barrameda, D., O’Conner, A.G., Barrows, J.M., Starr, A., Kovalik, T., Rabichow, B.E., Lehmkuhl, E.M., Shreiner, D.D., et al. (2019). Glycolysis upregulation is neuroprotective as a compensatory mechanism in ALS. Elife 8. 10.7554/eLife.45114.

Mariano, V., Achsel, T., Bagni, C., and Kanellopoulos, A.K. (2020). Modelling Learning and Memory in Drosophila to Understand Intellectual Disabilities. Neuroscience 445, 12–30. 10.1016/j.neuroscience.2020.07.034.

McAleese, K.E., Walker, L., Erskine, D., Thomas, A.J., McKeith, I.G., and Attems, J. (2017). TDP-43 pathology in Alzheimer’s disease, dementia with Lewy bodies and ageing. Brain Pathol 27, 472–479. 10.1111/bpa.12424.

McCarter, S.J., St Louis, E.K., and Boeve, B.F. (2016). Sleep Disturbances in Frontotemporal Dementia. Curr Neurol Neurosci Rep 16, 85. 10.1007/s11910-016-0680-3.

McDonald, K.K., Aulas, A., Destroismaisons, L., Pickles, S., Beleac, E., Camu, W., Rouleau, G.A., and Vande Velde, C. (2011). TAR DNA-binding protein 43 (TDP-43) regulates stress granule dynamics via differential regulation of G3BP and TIA-1. Human Molecular Genetics 20, 1400–1410. 10.1093/hmg/ddr021.

Meneses, A., Koga, S., O’Leary, J., Dickson, D.W., Bu, G., and Zhao, N. (2021). TDP-43 Pathology in Alzheimer’s Disease. Mol Neurodegener 16, 84. 10.1186/s13024-021-00503-x.

Mershin, A., Pavlopoulos, E., Fitch, O., Braden, B.C., Nanopoulos, D.V., and Skoulakis, E.M. (2004). Learning and memory deficits upon TAU accumulation in Drosophila mushroom body neurons. Learn Mem 11, 277–287. 10.1101/lm.70804.

Morante, J., and Desplan, C. (2008). The color-vision circuit in the medulla of Drosophila. Curr Biol 18, 553–565. 10.1016/j.cub.2008.02.075.

Muqit, M.M., and Feany, M.B. (2002). Modelling neurodegenerative diseases in Drosophila: a fruitful approach? Nat Rev Neurosci 3, 237–243. 10.1038/nrn751.

Murley, A.G., and Rowe, J.B. (2018). Neurotransmitter deficits from frontotemporal lobar degeneration. Brain 141, 1263–1285. 10.1093/brain/awx327.

Murphy, S.L., Kochanek, K.D., Xu, J., and Aria, E. (2021). Mortality in the United States, 2020. U.S. Department of Health and Human Services. https://www.cdc.gov/nchs/data/databriefs/db427.pdf.

Nana, A.L., Sidhu, M., Gaus, S.E., Hwang, J.L., Li, L., Park, Y., Kim, E.J., Pasquini, L., Allen, I.E., Rankin, K.P., et al. (2019). Neurons selectively targeted in frontotemporal dementia reveal early stage TDP-43 pathobiology. Acta Neuropathol 137, 27–46. 10.1007/s00401-018-1942-8.

Neumann, M., Sampathu, D.M., Kwong, L.K., Truax, A.C., Micsenyi, M.C., Chou, T.T., Bruce, J., Schuck, T., Grossman, M., Clark, C.M., et al. (2006). Ubiquitinated TDP-43 in frontotemporal lobar degeneration and amyotrophic lateral sclerosis. Science 314, 130–133. 10.1126/science.1134108.

Ostrowski, D., Kahsai, L., Kramer, E.F., Knutson, P., and Zars, T. (2015). Place memory retention in Drosophila. Neurobiol Learn Mem 123, 217–224. 10.1016/j.nlm.2015.06.015.

Park, J.H., Chung, C.G., Park, S.S., Lee, D., Kim, K.M., Jeong, Y., Kim, E.S., Cho, J.H., Jeon, Y.-M., Shen, C.K.J., et al. (2020). Cytosolic calcium regulates cytoplasmic accumulation of TDP-43 through Calpain-A and Importin α3. eLife 9, e60132. 10.7554/eLife.60132.

Pennington, C., Hodges, J.R., and Hornberger, M. (2011). Neural correlates of episodic memory in behavioral variant frontotemporal dementia. J Alzheimers Dis 24, 261–268. 10.3233/JAD-2011-101668.

Perry, D.C., Brown, J.A., Possin, K.L., Datta, S., Trujillo, A., Radke, A., Karydas, A., Kornak, J., Sias, A.C., Rabinovici, G.D., et al. (2017). Clinicopathological correlations in behavioural variant frontotemporal dementia. Brain 140, 3329–3345. 10.1093/brain/awx254.

Polymenidou, M., Lagier-Tourenne, C., Hutt, K.R., Huelga, S.C., Moran, J., Liang, T.Y., Ling, S.- C., Sun, E., Wancewicz, E., Mazur, C., et al. (2011). Long pre-mRNA depletion and RNA missplicing contribute to neuronal vulnerability from loss of TDP-43. Nature Neuroscience 14, 459–468. 10.1038/nn.2779.

Poos, J.M., Jiskoot, L.C., Papma, J.M., van Swieten, J.C., and van den Berg, E. (2018). Meta-analytic Review of Memory Impairment in Behavioral Variant Frontotemporal Dementia. J Int Neuropsychol Soc 24, 593–605. 10.1017/S1355617718000115.

Ramaswami, M., Taylor, J.P., and Parker, R. (2013). Altered Ribostasis: RNA-Protein Granules in Degenerative Disorders. Cell 154, 727–736. 10.1016/j.cell.2013.07.038.

Rascovsky, K., Hodges, J.R., Knopman, D., Mendez, M.F., Kramer, J.H., Neuhaus, J., van Swieten, J.C., Seelaar, H., Dopper, E.G., Onyike, C.U., et al. (2011). Sensitivity of revised diagnostic criteria for the behavioural variant of frontotemporal dementia. Brain 134, 2456–2477. 10.1093/brain/awr179.

Rawson, J.M., Dimitroff, B., Johnson, K.G., Rawson, J.M., Ge, X., Van Vactor, D., and Selleck, S.B. (2005). The heparan sulfate proteoglycans Dally-like and Syndecan have distinct functions in axon guidance and visual-system assembly in Drosophila. Curr Biol 15, 833–838. 10.1016/j.cub.2005.03.039.

Rohn, T.T. (2008). Caspase-cleaved TAR DNA-binding protein-43 is a major pathological finding in Alzheimer’s disease. Brain Research 1228, 189–198. 10.1016/j.brainres.2008.06.094.

Romano, M., Feiguin, F., and Buratti, E. (2012). Drosophila Answers to TDP-43 Proteinopathies. J Amino Acids 2012, 356081. 10.1155/2012/356081.

Rutherford, N.J., Zhang, Y.J., Baker, M., Gass, J.M., Finch, N.A., Xu, Y.F., Stewart, H., Kelley, B.J., Kuntz, K., Crook, R.J., et al. (2008). Novel mutations in TARDBP (TDP-43) in patients with familial amyotrophic lateral sclerosis. PLoS Genet 4, e1000193. 10.1371/journal.pgen.1000193.

Sani, T.P., Bond, R.L., Marshall, C.R., Hardy, C.J.D., Russell, L.L., Moore, K.M., Slattery, C.F., Paterson, R.W., Woollacott, I.O.C., Wendi, I.P., et al. (2019). Sleep symptoms in syndromes of frontotemporal dementia and Alzheimer’s disease: A proof-of-principle behavioural study. eNeurologicalSci 17, 100212. 10.1016/j.ensci.2019.100212.

Santiago, J.A., Bottero, V., and Potashkin, J.A. (2020). Transcriptomic and Network Analysis Identifies Shared and Unique Pathways across Dementia Spectrum Disorders. Int J Mol Sci 21. 10.3390/ijms21062050.

Seeley, W.W. (2008). Selective functional, regional, and neuronal vulnerability in frontotemporal dementia. Curr Opin Neurol 21, 701–707. 10.1097/WCO.0b013e3283168e2d.

Shaw, P.J., Cirelli, C., Greenspan, R.J., and Tononi, G. (2000). Correlates of sleep and waking in Drosophila melanogaster. Science 287, 1834–1837. 10.1126/science.287.5459.1834.

Sherman, B.T., Hao, M., Qiu, J., Jiao, X., Baseler, M.W., Lane, H.C., Imamichi, T., and Chang, W. (2022). DAVID: a web server for functional enrichment analysis and functional annotation of gene lists (2021 update). Nucleic Acids Res 50, W216-221. 10.1093/nar/gkac194.

Sreedharan, J., Blair, I.P., Tripathi, V.B., Hu, X., Vance, C., Rogelj, B., Ackerley, S., Durnall, J.C., Williams, K.L., Buratti, E., et al. (2008). TDP-43 Mutations in Familial and Sporadic Amyotrophic Lateral Sclerosis. Science 319, 1668–1672. 10.1126/science.1154584.

Sreedharan, J., Neukomm, L.J., Brown, R.H., Jr., and Freeman, M.R. (2015). Age-Dependent TDP-43-Mediated Motor Neuron Degeneration Requires GSK3, hat-trick, and xmas-2. Curr Biol 25, 2130–2136. 10.1016/j.cub.2015.06.045.

Stewart, B.A., Atwood, H.L., Renger, J.J., Wang, J., and Wu, C.F. (1994). Improved stability of Drosophila larval neuromuscular preparations in haemolymph-like physiological solutions. J Comp Physiol A 175, 179–191. 10.1007/BF00215114.

Stopford, C.L., Thompson, J.C., Neary, D., Richardson, A.M., and Snowden, J.S. (2012). Working memory, attention, and executive function in Alzheimer’s disease and frontotemporal dementia. Cortex 48, 429–446. 10.1016/j.cortex.2010.12.002.

Strausfeld, N.J., and Hirth, F. (2013). Deep homology of arthropod central complex and vertebrate basal ganglia. Science 340, 157–161. 10.1126/science.1231828.

Sun, Y., Qiu, R., Li, X., Cheng, Y., Gao, S., Kong, F., Liu, L., and Zhu, Y. (2020). Social attraction in Drosophila is regulated by the mushroom body and serotonergic system. Nat Commun 11, 5350. 10.1038/s41467-020-19102-3.

Susnjar, U., Skrabar, N., Brown, A.L., Abbassi, Y., Phatnani, H., Consortium, N.A., Cortese, A., Cereda, C., Bugiardini, E., Cardani, R., et al. (2022). Cell environment shapes TDP-43 function with implications in neuronal and muscle disease. Commun Biol 5, 314. 10.1038/s42003-022-03253-8.

Swain, A., Misulovin, Z., Pherson, M., Gause, M., Mihindukulasuriya, K., Rickels, R.A., Shilatifard, A., and Dorsett, D. (2016). Drosophila TDP-43 RNA-Binding Protein Facilitates Association of Sister Chromatid Cohesion Proteins with Genes, Enhancers and Polycomb Response Elements. PLoS Genet 12, e1006331. 10.1371/journal.pgen.1006331.

Team, R. (2022). RStudio: Integrated Development Environment for R. (PBC).

Team, R.C. (2021). A language and environment for statistical computing. (R Foundation for Statistical Computing).

Therneau, T.M. (2021). _A Package for Survival Analysis in R_. R package version 3.2–13.

Tollervey, J.R., Curk, T., Rogelj, B., Briese, M., Cereda, M., Kayikci, M., König, J., Hortobágyi, T., Nishimura, A.L., Župunski, V., et al. (2011). Characterizing the RNA targets and position-dependent splicing regulation by TDP-43. Nature Neuroscience 14, 452–458. 10.1038/nn.2778.

Tsao, C.H., Chen, C.C., Lin, C.H., Yang, H.Y., and Lin, S. (2018). Drosophila mushroom bodies integrate hunger and satiety signals to control innate food-seeking behavior. Elife 7. 10.7554/eLife.35264.

Vogt, K., Schnaitmann, C., Dylla, K.V., Knapek, S., Aso, Y., Rubin, G.M., and Tanimoto, H. (2014). Shared mushroom body circuits underlie visual and olfactory memories in Drosophila. Elife 3, e02395. 10.7554/eLife.02395.

Waghmare, I., Wang, X., and Page-McCaw, A. (2020). Dally-like protein sequesters multiple Wnt ligands in the Drosophila germarium. Dev Biol 464, 88–102. 10.1016/j.ydbio.2020.05.004.

Walker, A.K., Spiller, K.J., Ge, G., Zheng, A., Xu, Y., Zhou, M., Tripathy, K., Kwong, L.K., Trojanowski, J.Q., and Lee, V.M.Y. (2015). Functional recovery in new mouse models of ALS/FTLD after clearance of pathological cytoplasmic TDP-43. Acta Neuropathologica 130, 643–660. 10.1007/s00401-015-1460-x.

Wickham, H., Averick, M., Bryan, J., Chang, W., McGowan, L., François, R., Grolemund, G., Hayes, A., Henry, L., Hester, J., et al. (2019). Welcome to the Tidyverse. Journal of Open Source Software 4. 10.21105/joss.01686.

Wolff, G.H., and Strausfeld, N.J. (2016). Genealogical correspondence of a forebrain centre implies an executive brain in the protostome-deuterostome bilaterian ancestor. Philos Trans R Soc Lond B Biol Sci 371, 20150055. 10.1098/rstb.2015.0055.

Xiao, S., Sanelli, T., Dib, S., Sheps, D., Findlater, J., Bilbao, J., Keith, J., Zinman, L., Rogaeva, E., and Robertson, J. (2011). RNA targets of TDP-43 identified by UV-CLIP are deregulated in ALS. Mol Cell Neurosci 47, 167–180. 10.1016/j.mcn.2011.02.013.

Yamazaki, D., Horiuchi, J., Nakagami, Y., Nagano, S., Tamura, T., and Saitoe, M. (2007). The Drosophila DCO mutation suppresses age-related memory impairment without affecting lifespan. Nat Neurosci 10, 478–484. 10.1038/nn1863.

Yan, D., Wu, Y., Feng, Y., Lin, S.C., and Lin, X. (2009). The core protein of glypican Dally-like determines its biphasic activity in wingless morphogen signaling. Dev Cell 17, 470–481. 10.1016/j.devcel.2009.09.001.

Yu, H.H., and Lee, T. (2007). Neuronal temporal identity in post-embryonic Drosophila brain. Trends Neurosci 30, 520–526. 10.1016/j.tins.2007.07.003.

Zhan, L., Hanson, K.A., Kim, S.H., Tare, A., and Tibbetts, R.S. (2013). Identification of genetic modifiers of TDP-43 neurotoxicity in Drosophila. PLoS One 8, e57214. 10.1371/journal.pone.0057214.

Zhang, K., Guo, J.Z., Peng, Y., Xi, W., and Guo, A. (2007). Dopamine-mushroom body circuit regulates saliency-based decision-making in Drosophila. Science 316, 1901–1904. 10.1126/science.1137357.

Zwarts, L., Vanden Broeck, L., Cappuyns, E., Ayroles, J.F., Magwire, M.M., Vulsteke, V., Clements, J., Mackay, T.F., and Callaerts, P. (2015). The genetic basis of natural variation in mushroom body size in Drosophila melanogaster. Nat Commun 6, 10115. 10.1038/ncomms10115.

