## Supplemental Information for "Modelling dementia in *Drosophila* uncovers shared and specific targets of TDP-43 proteinopathy across ALS and FTD relevant circuits"

**AFFILIATIONS**:

Key Resources

| REAGENT or RESOURCE | SOURCE | IDENTIFIER |
|---|---|---|
| Antibodies | | |
| Anti-GFP-FITC | Rockland | Cat#: 600-402-215 |
| Anti-Fasciclin-II (ID4) | DSHB | RRID: AB_528235 |
| ChromoTek GFP-Trap® Magnetic Agarose | Proteintech | Cat#: gtma<br>RRID: AB_2631358 |
| Dally-like (13G8) | DSHB | RRID: AB_528191 |
| Polyclonal Alexa Fluor-568-conjugated goat anti-mouse secondary | ThermoFisher Scientific | Cat#A-11004;<br>RRID: AB_2534072 |
| Polyclonal Alexa Fluor-647-conjugated goat anti-mouse secondary | ThermoFisher Scientific | Cat#A-21235;<br>RRID: AB_2535804 |
| Living Color mouse Ab anti-GFP | Cell Signaling Technology | Cat# 2955 |
| Rabbit Ab beta-actin | Cell Signalling Technology | Cat#4967S |
| IRDye 800CW goat anti-mouse | LI-COR | Lot#D21115-25 |
| IRDye 680RD goat anti-rabbit | LI-COR | Lot# D21207-05 |
| Deposited data | | |
| Figure_1D_RatioAnalysis.csv | ScholarSphere | doi:10.26207/jq6p-w169 |
| Figure_1F_CellCountsAnalysis.csv | ScholarSphere | doi:10.26207/jq6p-w169 |
| Figure_2C_FluorescenceIntensity_RFP.csv | ScholarSphere | doi:10.26207/jq6p-w169 |
| Figure_2D_TDPYFPParticleSize.csv | ScholarSphere | doi:10.26207/jq6p-w169 |
| Figure_3_Y_maze.csv | ScholarSphere | doi:10.26207/jq6p-w169 |
| Figure_4A_4B_Day_Night_Sleep_SourceData.csv | ScholarSphere | doi:10.26207/jq6p-w169 |
| Figure_4C_SleepBoutLength_SourceData.csv | ScholarSphere | doi:10.26207/jq6p-w169 |

| | | |
|---|---|---|
| Figure_4C_SleepBoutNumber_SourceData.csv | ScholarSphere | doi:10.26207/jq6p-w169 |
| Figure_5_Survival_2022_SourceData.csv | ScholarSphere | doi:10.26207/jq6p-w169 |
| Figure_6A_Targets_SourceData.csv | ScholarSphere | doi:10.26207/jq6p-w169 |
| Figure_6B_6C_MB_MN_overlap_SourceData.csv | ScholarSphere | doi:10.26207/jq6p-w169 |
| Figure_7C_7D_DlpOE_YMazeSourceData.csv | ScholarSphere | doi:10.26207/jq6p-w169 |
| NCBI GEO Bioproject GSE217213 | NCBI | |
| Experimental models: Organisms/strains | | |
| *D. melanogaster*: -p65ADZp in attP40/ CyO; ZpGdbd in attP2 (MB-specific expression) | Yoshi Aso | *SS01276* |
| *D. melanogaster*: Oregon-R ; ; (OR-R, working memory assays) | Todd Schlenke | Oregon-R |
| *D. melanogaster*: OR-R; UAS-TDP-43$^{WT}$::YFP (working memory assays) | Shaun Davis (Schlenke laboratory), constructed from Estes et al., 2011; Estes et al.; 2013 | |
| *D. melanogaster*: $w^{1118}$ | Bloomington *Drosophila* Stock Center | RRID:BDSC_5905 |
| *D. melanogaster*: $w^{1118}$ ; UAS-TDP-43$^{WT}$ (untagged TDP-43 for sleep assays) | J. Paul Taylor (Ritson et al. 2020) | |
| *D. melanogaster*: $w^{1118}$ ; ; UAS-TDP-43$^{G298S}$ (untagged TDP-43 for sleep assays) | Takeshi Iwatsubo (Ihara et al. 2013) | |
| *D. melanogaster*: $w^{1118}$ ; UAS-TDP-43$^{WT}$::YFP; UAS mCD8::RFP (MB morphology and IPs) | Robert Kraft (Zarnescu laboratory) | |
| *D. melanogaster*: $w^{1118}$ ; UAS-TDP-43$^{G298S}$::YFP UAS-mCD8::RFP (MB morphology and IPs) | Robert Kraft (Zarnescu laboratory) | |
| $w^{1118}$ ; UAS-mCD8::RFP (control) | | BL 27391 |
| y $w^{1118}$ ; ; UAS-YFP (control) | | BL 6660 |

| Software | | |
|---|---|---|
| R | The R Foundation for Statistical Computing (2021) | 4.1.2 (2021-11-01) -- "Bird Hippie" |
| RStudio | RStudio Team (2021) | RStudio 2021.09.0+351 "Ghost Orchid" Release |

Tables

Summary tables in file Godfrey_et_al_Supplemental_Tables.xlsx provide summary statistics or interpetive data corresponding to the following figures and the corresponding supplemental:

| | |
|---|---|
| Table_S1E_NucCellRatio | Figure 1E |
| Table_S1F_CellNumber | Figure 1F |
| Table_S2A_mCD8RFP_Intensity | Figure 2A |
| Table_S2B_YFP_ParticleSize | Figure 2B |
| Table_S3_Ymaze | Figure 3 |
| Table_S4A_DayNightSleep | Figure 4A |
| Table_S4B_SleepBoutLength | Figure 4B |
| Table S4C_SleepBoutNum | Figure 4C |
| Table_S6A_WT_MBOnly_GO_Term | Figure 6A |
| Table_6B_G298S_MBOnly_GO_Term | Figure 6B |
| Table_S7C_S7D_DlpYmaze | Figure 7C and 7D |

Supplemental Figures

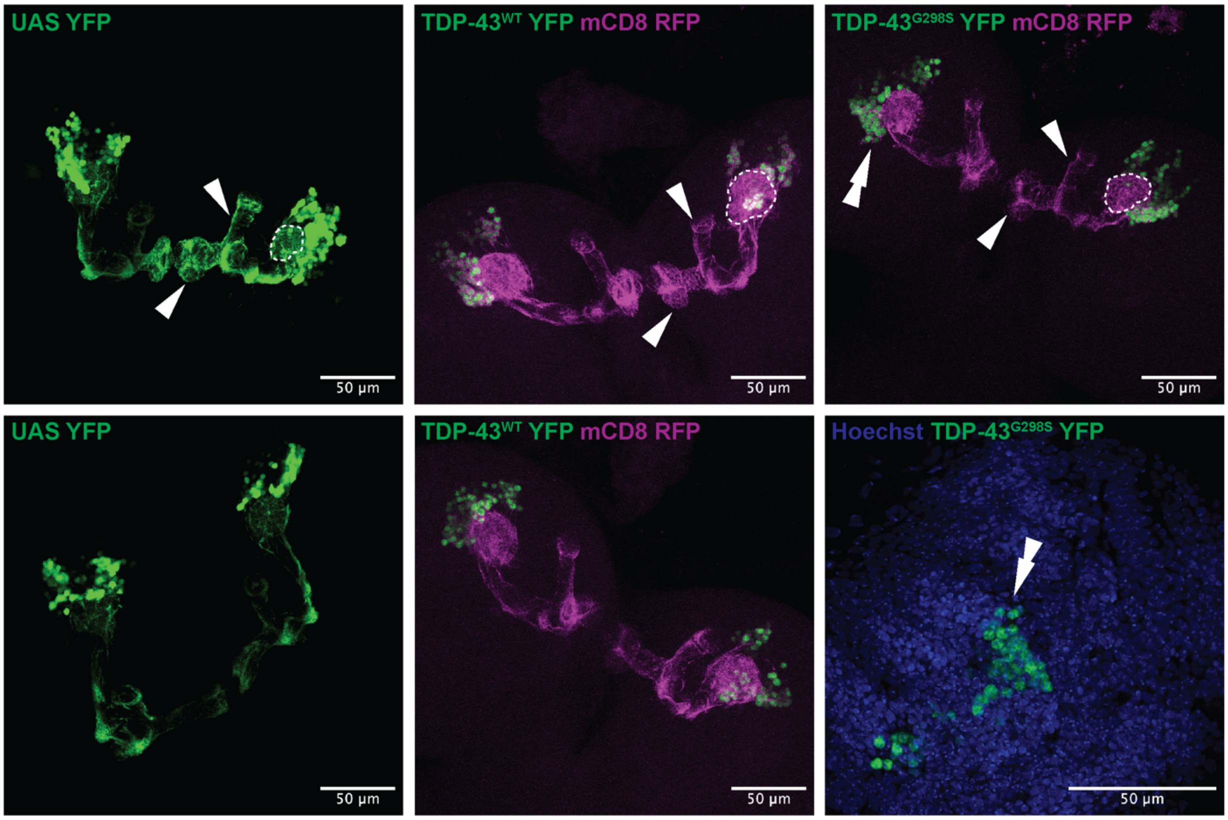


Figure 1-supplement 1. Maximum intensity projections showing split-GAL4 driver line *SS01276* expression in 3rd instar larva. YFP (left) labels cell bodies, calyx (dendrites, dotted outline), and lobes (axons, arrows). In TDP-43 YFP mCD8 RFP brains, TDP-43 YFP is restricted to cell bodies (middle and right) while mCD8-RFP allows visualization of all cell membranes. Measurement of TDP-43 YFP nucleus to whole cell ratio was performed from high magnification (100X) images of the calyx region (example lower left with double arrows corresponding to region in lower magnification image above).

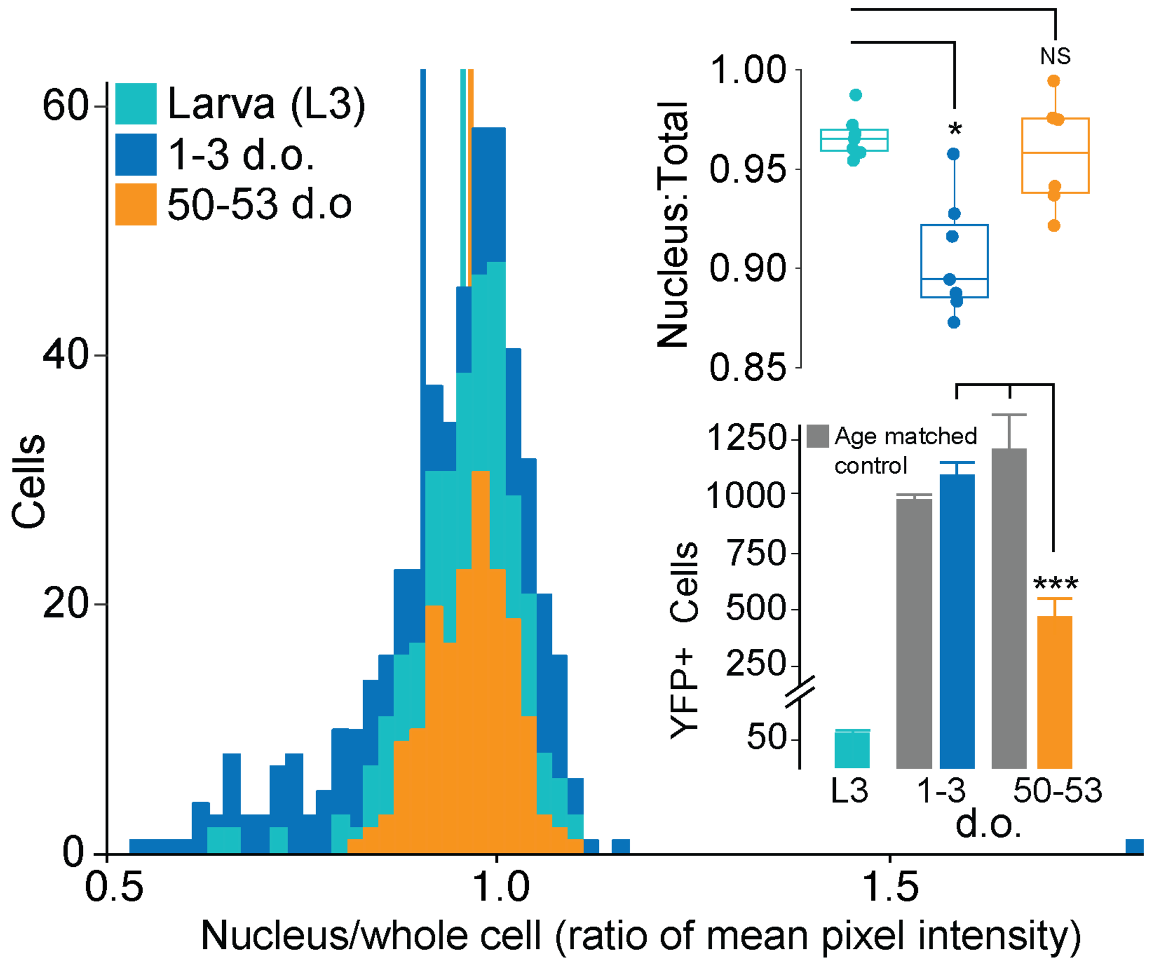


Figure 1-supplement 2. TDP-43$^{G298S}$ localization in MBN cell bodies. Histograms showing distribution of TDP-43 signal intensity ratios (cell nuclei to total cell) shift with age (days old, d.o.). Boxplot depicting age-specific TDP-43$^{G298S}$ signal intensity ratios. * = P < 0.05

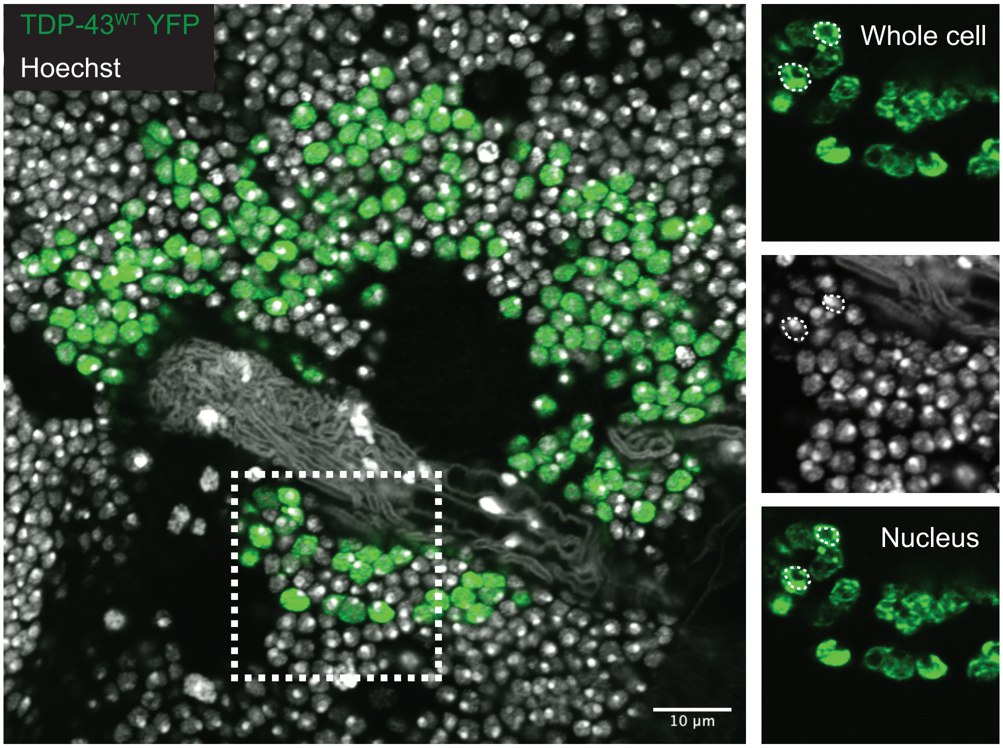


Figure 1-supplement 3. Method for measuring the ratio of nucleus to total cell TDP-43 YFP from mean pixel intensity of MBNs. White box on left depicts area of insets on right. Top, right inset shows cell body perimeter traced and measured in TDP-43 YFP channel (= total cell mean pixel intensity), middle and lower right panel show nuclear boundary traced in Hoechst channel and measured in TDP-43 YFP channel (= nucleus mean pixel intensity). Ratio of nucleus to total cell used for quantification of nuclear depletion in MBNs with age.

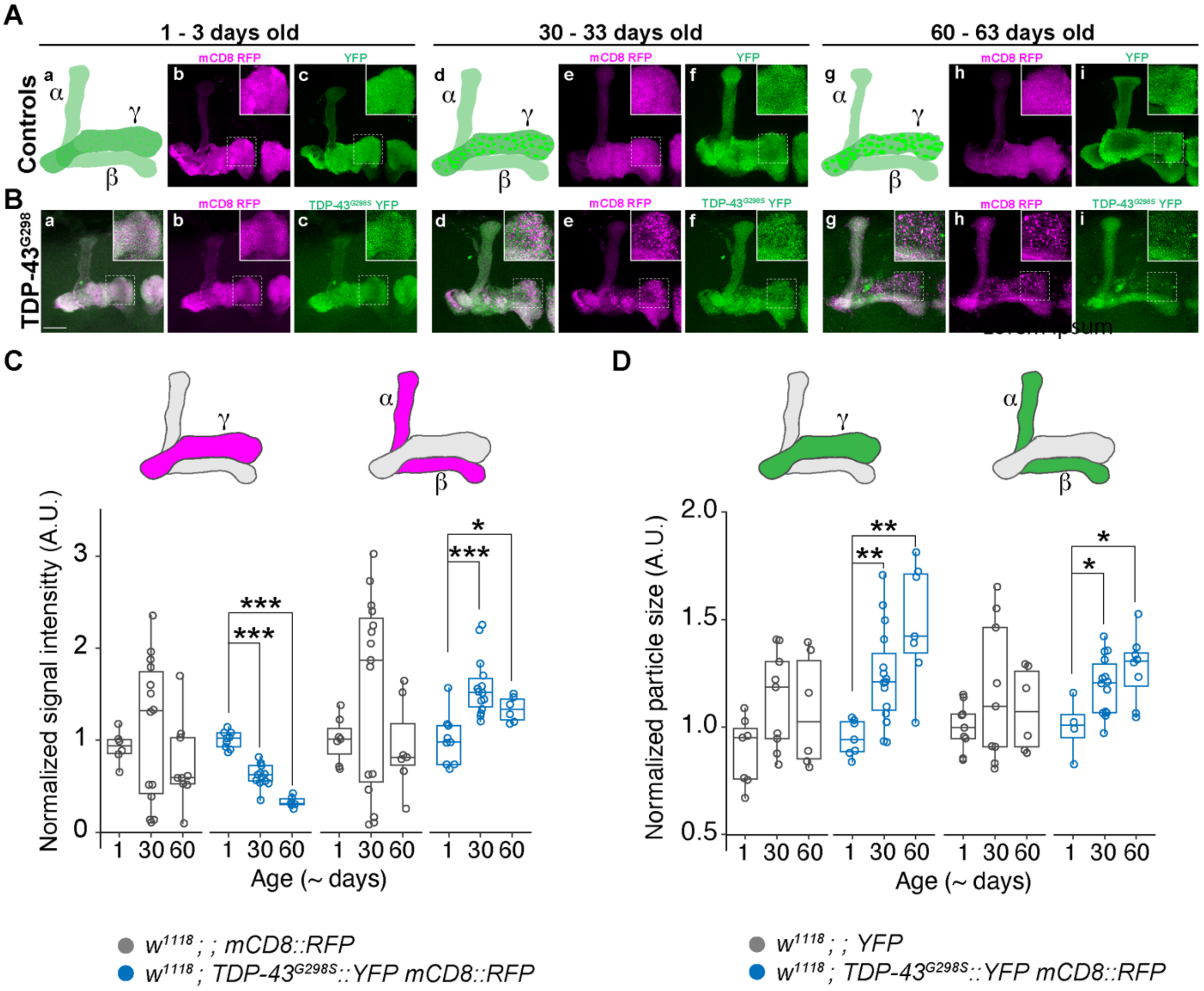


Figure 2-supplement 1. Mushroom body lobes (MBLs) show age-related, region-specific TDP-$43^{G298S}$ cytoplasmic accumulation and axonal fragmentation. (A) Illustrations of changes in MBLs targeted by TDP-43 OE over time (a, d, g) alongside morphology of young (b, c,) middle-aged (e, f) and old (h, i) control flies expressing membrane-bound RFP or cytoplasmic YFP. (B) Overexpression of TDP-$43^{G298S}$ in MBNs results in axonal localization of TDP-43 in young adult flies (c) and dystrophic neurites in middle-aged (e) and old (h) flies. (C) Signal intensity of RFP with age in $\gamma$ and $\alpha/\beta$ lobes. (D) Changes in YFP particle size with age in $\gamma$ and $\alpha/\beta$ lobes. * = P <0.05; ** = P < 0.01; *** = P < 0.001

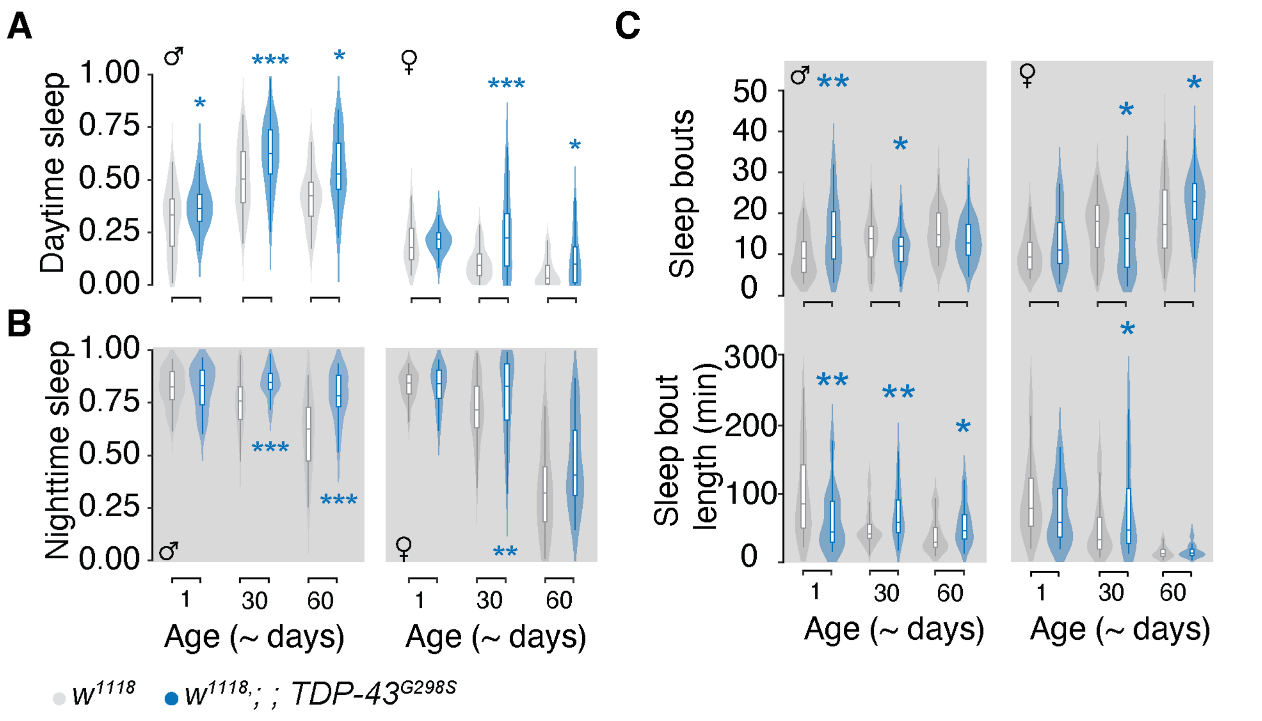


Figure 4-supplement 1. TDP-43$^{G298S}$ overexpression in MBNs reduces arousal, increasing day and night sleep. (A) Proportion of time flies spent sleeping during the day and (B) at night. (C) Sleep fragmentation assessed by number of sleep bouts (top panel) and mean bout length (bottom panel) during the night. Male data on the left and female data on the right in each panel. * = P <0.05; ** = P < 0.01; *** = P < 0.001

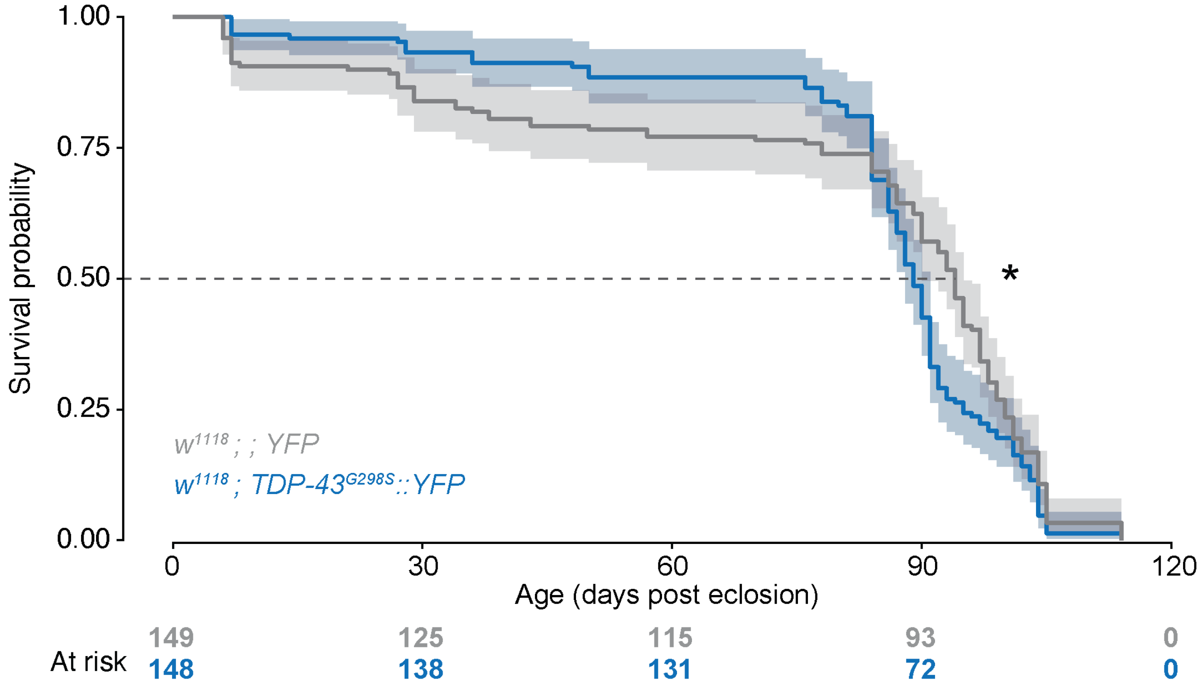


Figure 5-supplement 1. Mutant TDP-43 overexpression in MBNs is sufficient to reduce lifespan.

Data from male and female flies pooled for analysis. * P < 0.05

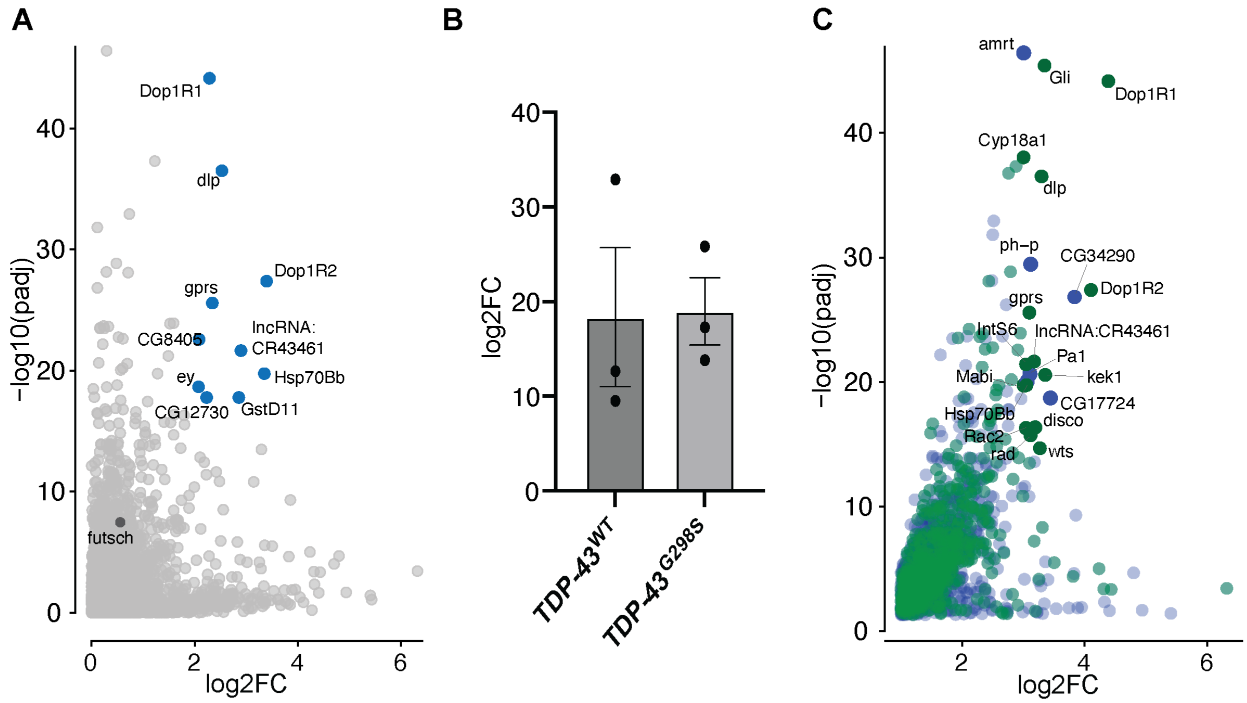


Figure 6-supplement 1. mRNAs enriched with TDP-43$^{G298S}$ overexpression in a *Drosophila* MBs. (A) Volcano plot displaying enriched mRNAs. Y-axis depicts Log2 Fold Change after subtraction of YFP control values. Blue circles indicate log2 fold change > 2 and P < 1 x $10^{-14}$. (B) qPCR validation of *dlp* enrichment in TDP-43 complexes immunoprecipitated from adult heads expressing TDP-43 with *SS01276*. (C) Volcano plot displaying mRNAs enriched in TDP-43$^{G298S}$ complexes that are MB-specific (blue) or shared between MB and MN models; saturated green circles indicate shared targets that show log2 Fold Change > 2 and P < 1 x $10^{-14}$; blue circles indicate MB targets with log2 fold change > 3 and P < 1 x $10^{-14}$.

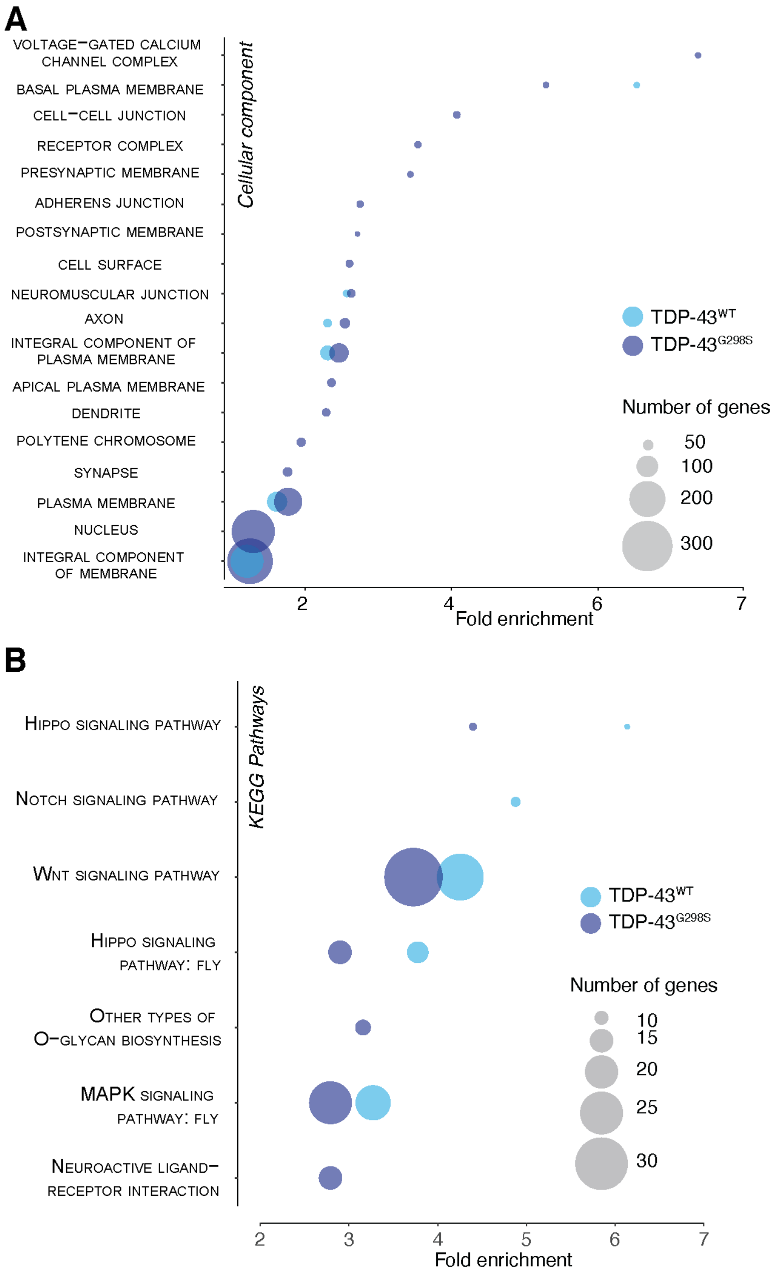


Figure 6-supplement 2. Functional annotation of enriched targets in fly models of TDP-43 driven dementia. (A) Cellular component of genes enriched in each TDP-43 model ranked by fold enrichment over all identified mRNAs in fly heads. (B) KEGG pathway analysis of targets enriched in each TDP-43 model ranked by fold enrichment over all identified mRNAs in fly heads.

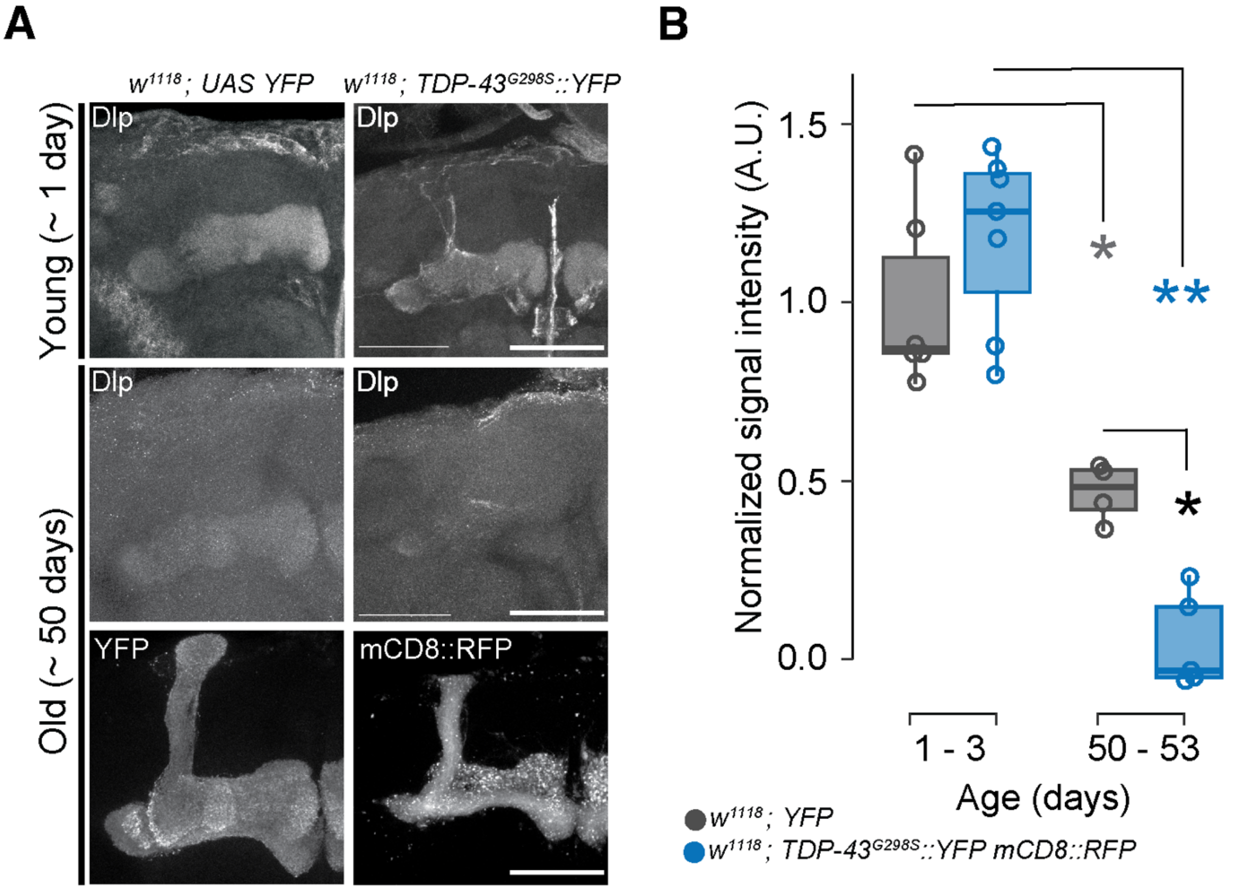


Figure 7-supplement 1. Dally-like protein is a target of TDP-43$^{G298S}$ in MBNs where it mediates TDP-43 dependent working memory deficits.. Deficits in working memory are evident in young adults. (A) Age-dependent loss of Dlp antibody labelling in mushroom bodies. Cytoplasmic YFP (Control) or mCD8 RFP (*TDP-43$^{G298S}$*) in aged flies indicating intact MBLs show decreased Dlp signal at ~ 50 days. Colored asterisks indicate statistical comparisons by age within a genotype. (B) Change in Dlp signal intensity in MBLs with age. Scale bar = 50 μm. * = P < 0.05, ** = P < 0 .01

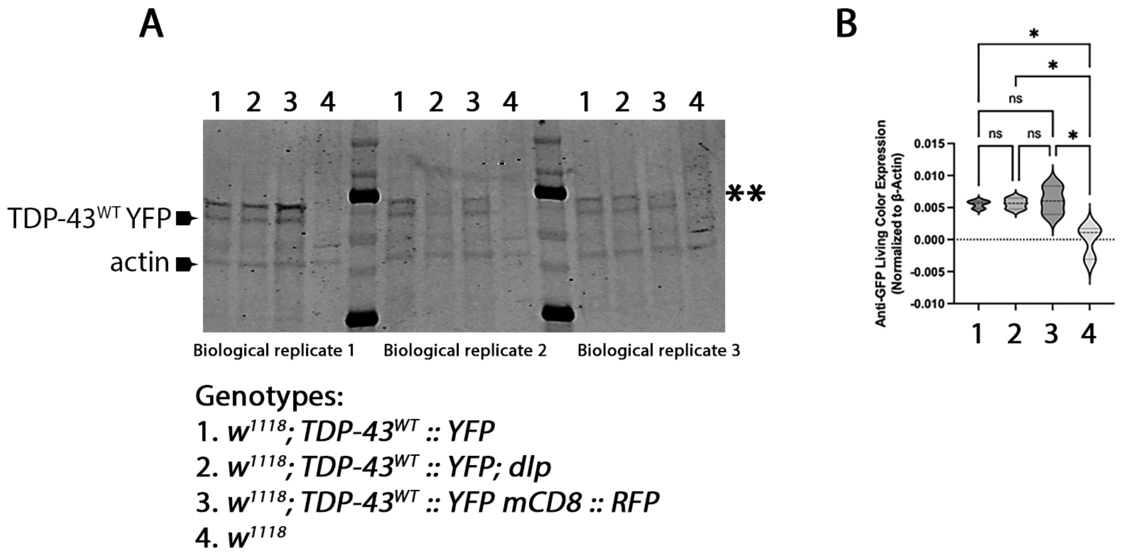


Figure 7-supplement 2. TDP-43$^{WT}$ protein expression is not reduced by the presence of a second UAS-driven transgene. (A) Western blot showing TDP$^{WT}$-43 YFP expression driven by the mushroom body driver line, *SS01276* in three different genotypes: *TDP-43$^{WT}$::YFP*, *TDP-43$^{WT}$::YFP dlp OE*, and *TDP-43$^{WT}$::YFP mCD8::RFP*. *w$^{1118}$* controls were used to confirm antibody specificity against GFP/YFP in fly tissues. TDP-43 YFP and actin, as indicated by arrows, left side. Double asterisk, right side, indicates non-specific band. (B) Quantification of western blot in (A). Genotypes and replicates, as indicated. * = P <0.05

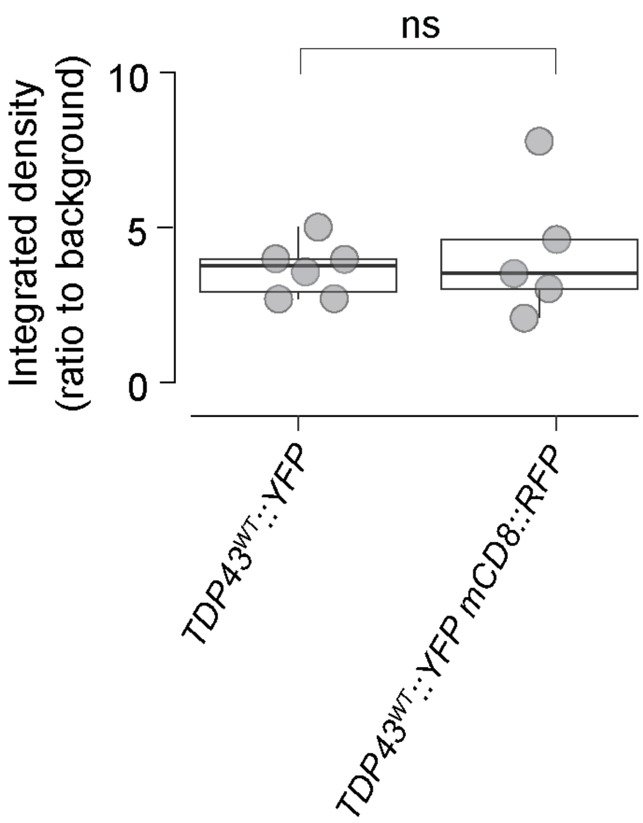


Figure 7-supplement 3. TDP-43$^{WT}$ YFP expression levels upon concommitant expression of a second UAS-driven transgene (mCD8 RFP). TDP$^{WT}$-43 YFP expression driven by the split-GAL4 mushroom body driver line, *SS01276* in two genotypes: *OR; TDP-43$^{WT}$::YFP* and *w$^{1118}$; TDP-43$^{WT}$::YFP mCD8::RFP*. Means compared using Wilcoxon Rank Sum.
